# The Darwinian evolution of political preferences and the collective strategies they support

**DOI:** 10.64898/2026.07.31.742050

**Authors:** Cedric Perret

**Affiliations:** University of Lausanne, Faculty of Business and Economics, Lausanne, Vaud, Switzerland

## Abstract

The success of social groups depends critically on the institutions, rules, and other collective strategies they adopt. Like material technologies, these social technologies may improve through a process analogous to biological evolution: if the preferences of more successful individuals are more likely to spread, groups may gradually adopt more effective strategies. Crucially, this could explain how complex yet efficient rules emerge. Still, it remains unclear whether this process generally generates improvement in collective strategies, and under what conditions. We address this question with a general model in which preferences over strategies are transmitted between individuals, while the strategies groups adopt are determined by those preferences. We show that selection on preferences can indeed improve collective strategies across a wide range of conditions. However, whether improvement occurs, and how quickly, depends strongly on how collective decisions translate individual preferences into group outcomes. We demonstrate that this can lead to dramatic long-run differences in the strategies groups adopt: some populations rapidly converge on efficient strategies, whereas others remain stuck with inefficient ones because improvement is too slow, stalls, or is undermined by drift or maladaptive biases.

## 1 Introduction

Social groups routinely adopt *collective strategies* that govern how members should act in a given situation: hunting bands agree on how to share a kill, governments set tax rates, and sports teams decide on their tactics. In fact, the very behaviours that researchers in social evolution typically focus on, how much to contribute to a public good or how severely to punish defectors, are often dictated by formal rules such as tax rates and standardised fines. The strategies that groups adopt are central to their success: they are widely recognised as a key determinant of societies’ economic performance (North, 1990; Acemoglu and Robinson, 2012), the primary means by which communities manage common resources (Ostrom, 1990), and a key mechanism that enabled the emergence of large-scale societies (Powers et al., 2016).

As such, understanding how such strategies change over time is a central goal across the social and evolutionary sciences. Here we examine the recurring but understudied idea that change in collective strategies could be driven by a selective process analogous to biological evolution (Alchian, 1950; Boyd and Richerson, 2009; Hannan and Freeman, 1984; Smaldino, 2014; Molho et al., 2024). Specifically, we examine the idea that collective strategies improve because people tend to adopt their preferences over which strategies to support from more successful individuals.

This is neither a far-fetched idea nor a trivial analogy. Humans have been shown to learn from successful individuals across a range of domains (Richerson and Boyd, 2004; Henrich and Henrich, 2007; Burton-Chellew and D’Amico, 2021). Furthermore, political preferences are partly transmitted from parents to offspring within families (Jennings et al., 2009; Durmuşoğlu et al., 2023; Dawes and Weinschenk, 2020), which would likewise spread the preferences of more successful individuals, since such individuals tend to leave more offspring. In line with these, several existing models already assume that political preferences change through such processes when examining the emergence of specific collective strategies (Lehmann and Feldman, 2008; Powers and Lehmann, 2013; Johnstone et al., 2020; Mullon and Lehmann, 2022; Powers et al., 2023; Hunt et al., 2024).

Furthermore, this process provides a route by which collective strategies would improve even when nobody has a causal understanding of how or why they work (Derex et al., 2019). This would offer a parsimonious explanation for how complex but efficient rules and institutions can emerge, or fail to emerge. These situations may often occur in practice since evaluating the consequences of collective strategies requires accounting for many complex and indirect interactions. In fact, evidence indicates that individuals often lack the information or ability to determine the consequences of alternative rules (Bartels, 2005; Gemmell et al., 2004), or ignore such considerations altogether (Delli Carpini and Keeter, 1997; Campbell et al., 1960). Yet these implications hold only if the proposed mechanism does in fact improve collective strategies. Whether it does so remains unclear.

If anything can be learned from past attempts to draw parallels between biological and cultural change (Lewens, 2015; Claidière et al., 2014), it is that conclusions from evolutionary theory cannot be carried over to other domains without caution. Ideas are not genes. And as ideas are not genes, political preferences are not “just ideas”. In particular, whether copying improves a trait depends on traits associated with greater success being more likely to be copied (Boyd and Richerson, 1985), an association that for instance can be weakened by various learning biases (Henrich and McElreath, 2003; Aoki et al., 2011). Yet, a distinctive feature of political preferences is that they affect success only indirectly, through the collective strategies they help produce. Because any one individual may have limited influence over the strategy their group adopts, their success may only be weakly related, or not related at all to their preferences. For instance, someone may prosper yet favour a poorly designed collective strategy because they belong to a group that adopts a better one. Thus, copying these preferences need not spread the ones that generate better strategies.

So far, we know that learning political preferences from successful individuals can improve collective strategies in some cases because models using this process to study particular strategies show such improvement (Lehmann and Feldman, 2008; Powers and Lehmann, 2013; Mullon and Lehmann, 2022; Powers et al., 2023; Hunt et al., 2024). Yet, these examples are limited, typically assuming only one or two ways for groups to decide on their strategies, and, as explained above, there are clear reasons to expect the process to fail under some conditions. The remaining question, then, is whether the spread of political preferences from successful individuals generally generates cumulative improvement in the strategies groups adopt over time. Can this process be limited or fail, and if so, under what conditions? Answering this question would yield the kind of predictions that have made evolutionary reasoning useful for understanding cultural change (Kline and Boyd, 2010; Rogers and Ehrlich, 2008; Mesoudi, 2011).

To address this gap, we develop a general evolutionary model in which political preferences are the evolving traits and collective strategies emerge through collective choice. Using this framework, we show that whether selection can improve these strategies, and how much they improve, can vary sharply across cases, and that this variation depends critically on how collective decisions aggregate individual preferences into collective strategies. Our study continues a long-standing programme of models in cultural evolution that asks when social transmission generates cumulative improvement in cultural traits (studying, for instance, how different modes and biases of social learning shape cultural evolution; Cavalli-Sforza and Feldman, 1981; Boyd and Richerson, 1985), here examining a different class of traits, as has been called for (Smaldino, 2014; Molho et al., 2024).

## 2 A model of Darwinian evolution of political preferences

### 2.1 Conceptual framework

A collective strategy is a collective decision affecting how group members behave in a given situation (see Appendix A for our terminology). It either (i) prescribes precise actions members will take (e.g. “contribute 30% of your income to a public good”), or (ii) modifies the set or payoffs of available actions (e.g. “do not catch more than 30 fish per day”). A political preference is an individual’s preferences over alternative collective strategies (e.g. “everybody should contribute 30% of their income to the public good”). The preferences of individuals in a group jointly determine the collective strategy the group adopts, but the preference of *any* given individual may or may not affect the implemented strategy.

A cultural model of Darwinian evolution of political preferences is one in which the distribution of political preferences changes over time because preferences are transmitted through social learning, and those associated with higher payoffs are more likely copied. These models typically assume a population structured into groups repeatedly facing a strategic situation that generates material payoffs (the economic game). Actions in this game are determined by the collective strategy, itself determined by the preferences of potentially all group members (the political game) (Hurwicz, 1996; Powers et al., 2023; Gavrilets and Currie, 2023). This contrasts with standard evolutionary game theory, where an individual’s action depends solely on its trait (Figure 1).

**Figure 1.**
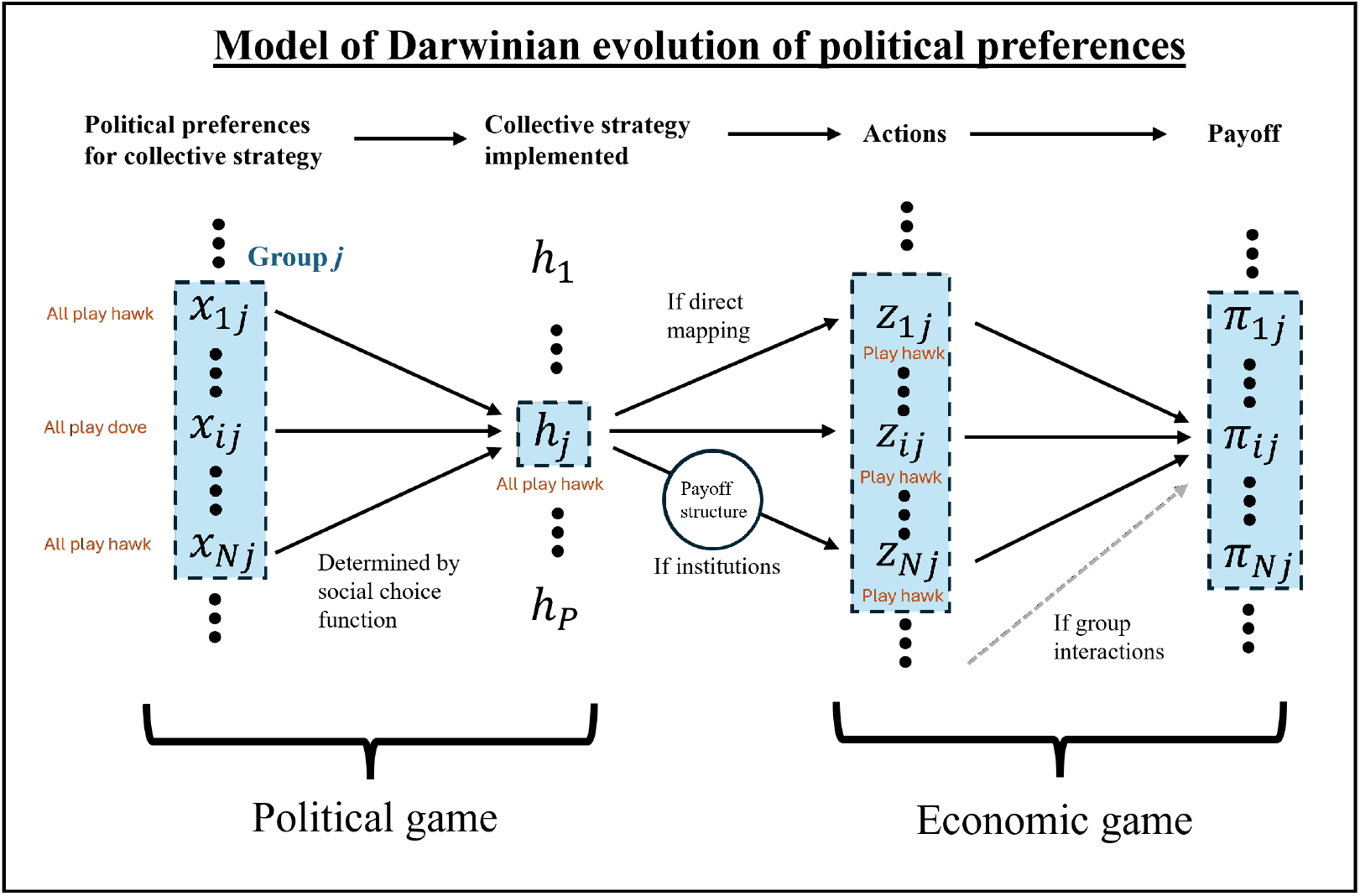
Generic representation of models of evolution of political preferences (Hurwicz, 1996; Powers et al., 2023; Gavrilets and Currie, 2023). Political preferences determine the collective strategy via a social choice function; the collective strategy then shapes actions in a recurrent economic game, generating payoffs that affect the transmission of preferences. Unlike standard evolutionary game theory, the transmitted trait is a preference over the actions of all group members rather than directly the carrier’s action. For example, in standard models of punishment, the trait sets each individual’s investment in punishment; in a collective strategy model, traits determine everyone’s investment in punishment.

We study a deliberately simple model in which we only assume that new political preferences arise through random variation and successful ones spread more widely. We do not model the mechanisms of social learning in detail, other forces affecting transmission and any process by which individuals might revise their preferences. This model is the closest analogue to biological evolution and provides a benchmark for how much improvement selection alone can generate. We relax some of these assumptions in Section 4.2.

### 2.2 Model setup and life cycle

We consider a population structured in *M* groups of constant size *N*. We index groups by *j* = 1, …, *M* and individuals within group by *i* = 1, …, *N*. Each group *j* at time *t* has a collective strategy, *h*_*j*_(*t*) ∈ [0, 1]. We assume that it varies continuously along a single dimension between two extremes represented by 0 and 1 (e.g. none to full contribution). An individual *i* in group *j* carries a political preference *x*_*ij*_(*t*) ∈ [0, 1], defined on the same domain as collective strategies (following the assumptions in Section 2.2.1).

At each time step, individuals undergo the following discrete-time life cycle: (i) collective strategies *h*_*j*_(*t*) are set based on the political preferences *x*_*ij*_(*t*) of group members; (ii) collective strategies affect the actions that individuals take in a subsequent, unspecified economic game, which in turn determine their payoffs *π*_*ij*_(*t*); (iii) a new population of political preferences *x*_*ij*_(*t* + 1) is formed by sampling preferences of individuals from the whole population with probabilities proportional to their payoffs.

We now describe each step of the life cycle in more detail.

#### 2.2.1 Collective decision-making

Collective strategies are set through a (collective) decision-making process, in which group members express their opinions, debate and/or vote, until they agree on a single decision. This process can take many forms but as formalised by social choice theory (Arrow, 1951; Moulin, 1988), any collective decision process can be summarised by a function that maps members’ political preferences 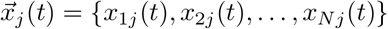 to the chosen collective strategy *h*_*j*_(*t*)

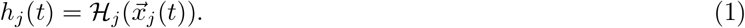

This function is called a social choice function (SCF). Examples of SCFs are a function that returns the average preference in the group or one returning the preference of a single individual; we compare such cases below. Importantly, the SCF summarises, but does not describe the concrete decision procedure (e.g. who speaks, how votes are cast, or how agreement is reached). Indeed, different procedures can be summarised by the same SCF: average may arise either from a voting system where the mean of all votes is chosen, or from informal discussions among individuals with similar status, speaking time, and influence. Conversely, similar procedures can correspond to different SCFs when influence or status differs across individuals.

A general form of political preferences would be a utility value that individuals assign to each possible collective strategy. However, we restrict attention to preferences with a simple representation. First, we assume that each individual has a *single* most preferred value which they would choose over all the others (Black, 1948), and second, that the SCF determines the collective strategy solely from these most preferred values. These imply that each individual’s preference can be summarised by a single value, which is the strategy they would set if they were the sole decision-maker. This makes preferences more tractable as evolving traits and matches how they are modelled in evolutionary models (Lehmann and Feldman, 2008; Mullon and Lehmann, 2022; Powers et al., 2023; Hunt et al., 2024). Finally, we consider reasonable SCFs that satisfy the property of unanimity, so that if everyone has the same preferred strategy, the function selects it.

#### 2.2.2 Payoff

Once collective strategies are established, individuals take part in an economic game that sets their payoffs. As a first step, we assume that all individuals in a group obtain the same payoff (we come back to this assumption in the discussion). We first write the payoff *π*_*ij*_(*t*) *>* 0 of individual *i* in group *j* in a general form:

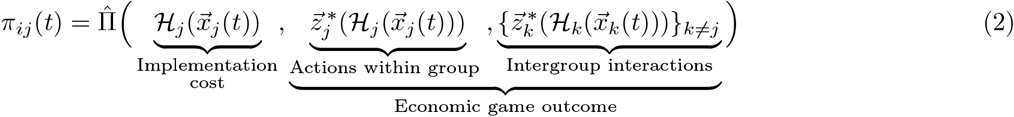

The payoff depends on (i) a direct cost of implementing the strategy and (ii) the outcome of the economic game, which is determined by the actions of individuals involved denoted 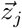 for members of focal group *j*, and possibly the actions of individuals in other groups, e.g. intergroup conflict. These actions may depend on the strategies in place either directly, when collective strategies prescribe behaviour, or indirectly, when collective strategies alter the strategic environment faced by individuals. In the latter case, actions are given by some equilibrium of the game induced by the collective strategies. We assume that behavioural adjustment occurs on a much faster timescale than the dynamics of political preferences, so that only this equilibrium behaviour matters.

For the purposes of our analysis, we do not specify the economic game and instead assume a general *payoff function* that maps each collective strategy to a non-negative payoff:

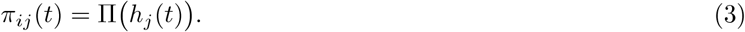

We present here the version without intergroup interactions, which is the one analysed in the main text; the full model is presented in Appendix B. This version is sufficient for our main question: given that some collective strategies yield higher payoffs than others—some hunting tactics perform better than others, some rules are better at maintaining cooperation—does selection on political preferences drive the strategies toward those values?

#### 2.2.3 Social learning

The key assumption of this model is that political preferences are transmitted in proportion to the payoffs obtained by their carriers. Concretely, at each time step, the *N* political preferences in each group (one per individual) are replaced by sampling, with replacement, from the pool of preferences in the whole population, with probabilities proportional to individual payoffs (Wright–Fisher process; Ewens, 1979). The probability that the political preference *x*_*ij*_(*t*) of individual *i* in group *j* is sampled in a given draw is

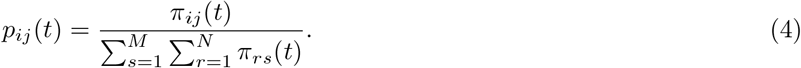

We assume that the probability that one political preference replaces another does not depend on whether their carriers belong to the same group (we relax this assumption in Appendix C). This procedure for forming the new population is standard and deliberately abstract. For intuition, it can be interpreted as the formalisation of two familiar models (Micheletti, 2020): (i) a biological model in which individuals reproduce asexually, have a number of offspring proportional to their payoffs and transmit their preferences vertically to their offspring, or (ii) a cultural model in which individuals update their preferences through horizontal, payoff-biased social learning.

To generate variation, we assume that transmission is occasionally imperfect. With probability *µ*_*m*_, the sampled political preference is replaced by a new value drawn from a Gaussian distribution centred on the sampled value, with standard deviation *σ*_*m*_ and truncated to [0, 1]. These changes occur in random directions, and individuals do not otherwise adjust their preferences.

### 2.3 Method for analysis

The previous section defines a generic model for the Darwinian evolution of political preferences. Once the economic game and the social choice function are specified, this framework can recover familiar models from the literature; we illustrate this later by recovering two such models (Case studies 1 and 2; Appendix E). Our aim is now to determine when collective strategies improve over time, how fast change occurs, and where it settles. To do so, we adopt a small-variance approximation to derive general expressions for these quantities and (ii) use individual-based simulations to assess how these predictions carry over to full stochastic dynamics in finite, more realistic populations.

#### 2.3.1 Analytical

To obtain analytical results, we make the standard assumptions that random modifications are rare and their effects are small, so that variation in political preferences is approximately normally distributed and maintained at low levels by a balance between stabilising selection and a constant influx of noise (Avila and Mullon, 2023). Under this approximation, we treat the average collective strategy as the strategy associated with the average political preference. We therefore track changes in the average preference, which is governed by the selection gradient

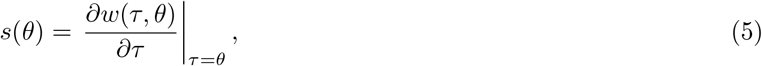

that is the effect of a marginal change in the preference *τ* of a focal individual away from the population mean *θ*, on its fitness *w*(*τ, θ*). For simplicity, we omit the time dependence (*t*) here and below. Long-term evolutionary outcomes are then characterised by the zeros of this gradient and by higher-order derivatives (see Appendix B for details).

The fitness of an individual is its expected number of copies over one time step. Substituting eqs. (1) and (3) into eq. (4), and setting the preference of individual *i* to *τ* and the preferences of all other individuals to *θ*, gives the fitness of the focal (eq. (A-6)). Assuming a large number of groups, so that the focal group has a negligible effect on the mean population fitness, the fitness of the focal is approximated by

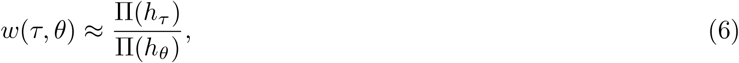

where *h*_*τ*_ = ℋ (*τ, θ*, …, *θ*) is the strategy of a group where one individual has preference *τ* and all others have preference *θ*, and *h*_*θ*_ = ℋ (*θ*, …, *θ*) is the strategy of a group where all individuals have preference *θ*.

#### 2.3.2 Simulations

Simulations follow the life cycle described in Section 2.2 and are implemented in a finite population structured into *M* groups of size *N*. To conduct simulations, we also specify the payoff function, starting with the simplest case in which the collective strategy has a single optimal value:

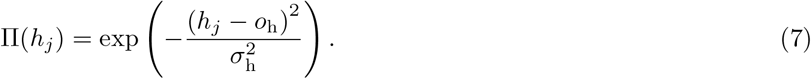

The parameter *o*_h_ ∈ [0, 1] is the strategy that yields the maximum payoff, and *σ*_h_ *>* 0 controls how sharply payoffs decline away from it. One example that can generate such a function is a collective strategy specifying a mandatory contribution (or tax) to a public good. If we assume that (i) the public good yields returns that exceed its cost, but that (ii) individuals may not accurately perceive that it is the case, then a higher tax rate initially raises total contributions and thus payoff, but excessively high taxation leads to tax avoidance and reduced incentives to produce (Wanniski, 1978). Later, we also consider a more complex payoff function and fully specified economic games as case studies; these are described in the corresponding sections. Specific SCFs are introduced where they are analysed.

## 3 How selection acts on political preferences and collective strategies

We now characterise how selection acts on political preferences, and what this implies for the evolution of collective strategies, in the case without intergroup interactions (see Appendix B for the complete analysis).

### Selection when strategies act as individual traits

We first establish a baseline by examining the case in which an individual preference maps directly to the implemented collective strategy and hence to its success. In our model, this baseline is obtained by setting group size *N* = 1. This recovers the case of the evolution of a standard trait. It also covers models (Isakov and Rand, 2012; Gavrilets and Duwal Shrestha, 2021) and theories (Boyd and Richerson, 1985, 2009) which assume implicitly that the collective strategies themselves are the traits being transmitted, and transmission occurs directly between groups (or between single individuals standing in for groups). In this case, the selection gradient is

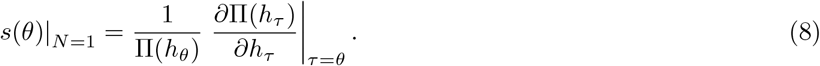

In this baseline, it coincides with the payoff gradient with respect to collective strategies, that is, the effect of a marginal change in the implemented strategy (and not the preference), on the payoff obtained in the focal’s group. The sign of this gradient gives the local direction in which collective strategies would yield higher payoff, and selection therefore moves preferences in that direction. We show in Appendix B that this takes place until the average preference reaches (i) a boundary of the feasible collective strategy space (*h* = 0 or *h* = 1) or (ii) a stable interior singular point *θ*^∗^ defined by

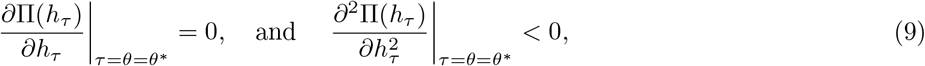

that is, a collective strategy that locally maximises payoff. We now ask how this changes once the implemented strategy is determined jointly by the preferences of several group members.

### Selection when strategies are collectively determined

For groups of any size *N*, the selection gradient becomes

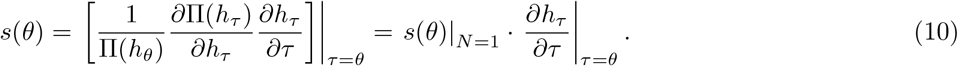

Equation (10) gives the central theoretical result of the paper. Selection on political preferences is driven by the same gradient as if strategies evolve directly, but multiplied by an additional term: the extent to which a marginal change in an individual’s political preference affects the implemented collective strategy. We call this term *marginal influence*.

If marginal influence is positive, the selection gradient has the same sign as the payoff gradient and thus, selection still favours changes in preferences towards strategies that yield higher payoff. Appendix B shows that it does so until the population reaches the same boundary or the same local maximum defined in (9) (branching can occur only in the case with group interactions; see Appendix B.6). Thus, given sufficient time and across a wide range of cases, the same locally optimal collective strategy evolves (the same as if collective strategies themselves evolve directly). The important difference is that the condition for this outcome, and the dynamics leading to it, are no longer the same.

First, improvement occurs only if the marginal influence is positive. This holds for SCFs with the property that, whenever one individual shifts their preference in one direction while others remain unchanged, the resulting collective strategy shifts in the same direction (a relaxed version of strong monotonicity conditions often discussed in social choice theory (Fishburn, 1973)). This also includes more general SCFs in which shifts in some individuals’ preferences have no effect, or even move the strategy in the opposite direction, provided these are outweighed by individuals with sufficiently positive influence.

Second, even under positive marginal influence, its magnitude determines the strength of the selection gradient, and consequently the rate of change. If marginal influence is too small, selection pressure is weak, and change may proceed too slowly for the population to approach the optimal collective strategy within a reasonable timescale. If it is sufficiently large, by contrast, change can proceed as quickly as if collective strategies themselves evolved directly.

Having characterised in general when selection improves collective strategies, we now return to our broader question of how reliably such improvement occurs. Our last result did not yet answer the question, because it is not straightforward what value marginal influence takes for a given SCF, or how much it varies across SCFs. We therefore now examine evolution under four specific SCFs.

## 4 How reliably does selection improve collective strategies?

### 4.1 From rapid to stalled improvement

#### Social choice functions considered

We now consider four specific SCFs: the collective strategy is the (i) *average* of preferences; (ii) *median* of preferences; (iii) the preference of a single *dictator* randomly selected once at the beginning of the simulation; or (iv) the preference with the greatest *novelty* within the group, defined for group *j* as

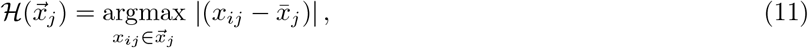

where 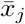 is the mean preference. The first three are standard SCFs in social choice theory (Moulin, 1988). The last one, novelty, is introduced to illustrate our main result. It may represent a society that favours novel ideas, either explicitly through how decisions are taken or because of individual biases.

We first derive predictions for these SCFs under the analytical approximation and then test in simulations how well they carry over to finite populations. Under the small-variance approximation, eq. (10) shows that differences in the rate of improvement are captured by the marginal influence 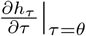. We obtain this term by evaluating the strategy *h*(*τ, θ*) of a group with one focal individual with preference *τ* and *N* − 1 with preferences *θ*.

#### Rate of improvement under average rule

Under the *average* rule,

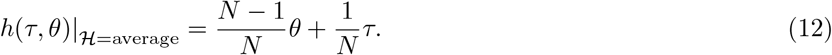

Substituting this equation in eq. (10), the selection gradient is

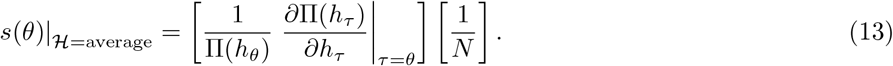

The direction of change is the same as the payoff gradient with respect to the collective strategies, but the rate of change drastically decreases as groups get larger. In other words, selection would improve collective strategies in this population (even if individuals randomly modify the rules) but only while groups remain small. In large groups, selection weakens, and change is potentially too slow for the groups to reach the optimal value in a reasonable time. This result does not require group size to directly hinder decision-making, for example through longer deliberation.

We test this prediction in Fig. 2 by varying *N* while keeping total population size constant. The simulations are consistent with the predictions: collective strategies improve under all *N*, but convergence is very slow for larger group sizes, leaving these groups far from the optimum even over long horizons. This slowdown translates into substantial payoff differences across populations: after 50,000 time steps, individuals in populations with group size *N* = 10 earn on average nearly five times the payoff of those with group size *N* = 100 (top-right panel, Figure 2).

**Figure 2.**
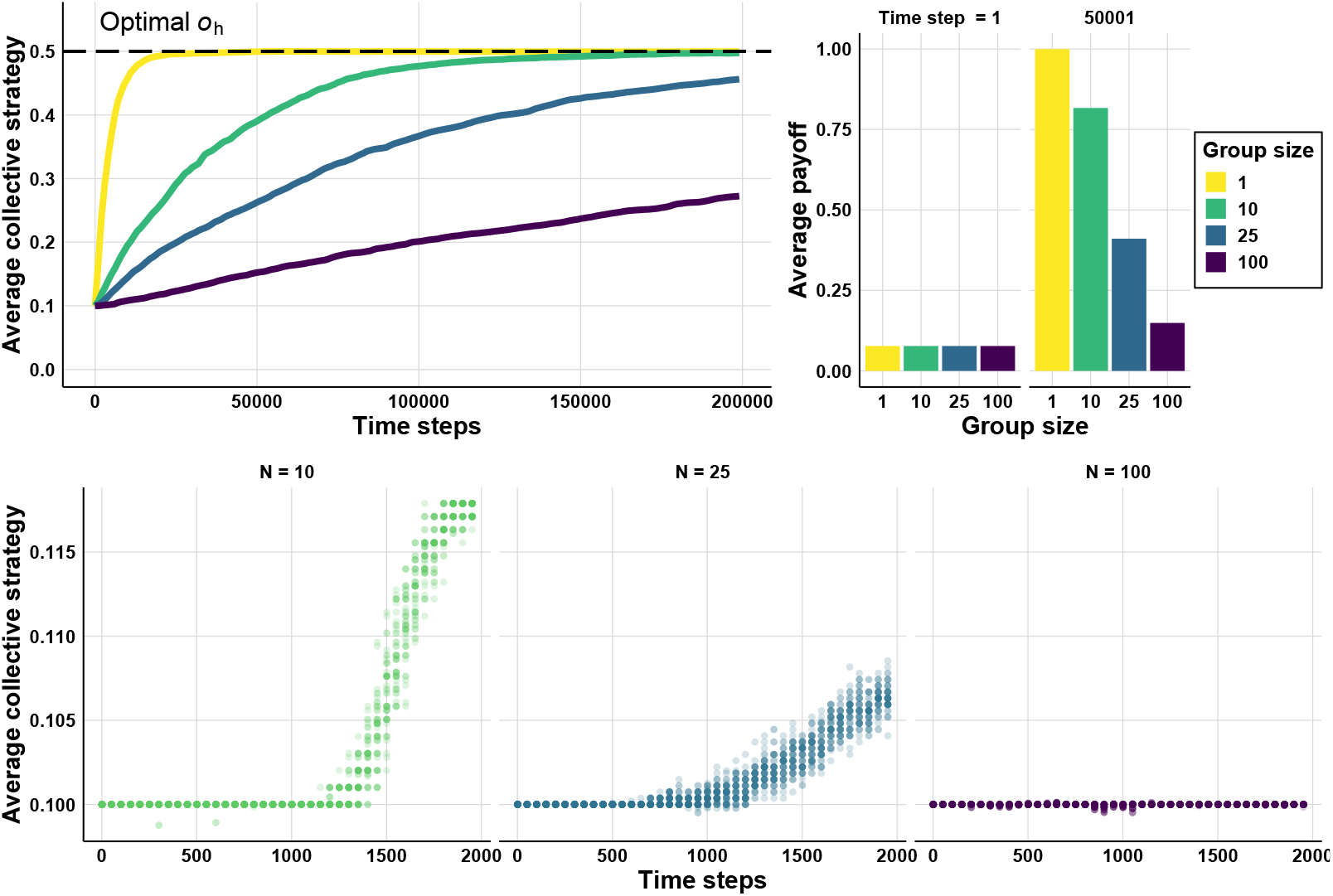
Evolution of collective strategies *h* over time for different group sizes when the strategy is the average of preferences. Top-left panel shows the mean 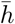 across 50 replicates, bottom panel shows *h*_*j*_ for 50 groups in a single run and top-right panel shows the mean payoff for different group sizes at time step 1 and 50001. We consider a large population of fixed size (10 000), partitioned into groups of size *N*, so that the number of groups is *M* = 10 000/*N*. Initial preferences are *x*(0) = 0.1. The payoff function has a single optimal value *o*_h_ = 0.5 and width *σ*_h_ = 0.25. The rate of random modifications is *µ*_m_ = 1 *×* 10^−5^, and the SD of random modifications is *σ*_m_ = 0.01.

#### Rate of improvement for different social choice functions

We then repeat the same calculation for the other SCFs (Table 1). Marginal influence varies widely across SCFs: it disappears entirely under the *median* rule (for *N* ≥ 3), preventing collective strategies from improving, while it is equal to 1 under *novelty*, leading to the same rate as in the baseline *N* = 1. Furthermore, the marginal influence equals 1/*N* under both *average* and *random dictator*, highlighting that rates can coincide across seemingly very different procedures. This happens because with *random dictator*, one individual has influence 1 and the others 0, so the expected influence of a random individual is still 1/*N*. A random individual either has a large effect rarely or no effect most of the time.

**Table 1.** Collective strategy in a group with a single focal individual, and the resulting marginal influence (which modulates the rate of change in political preferences).

| Social choice function | Collective strategy of<br>focal $h(\tau, \theta)$ | Marginal influence<br>$\frac{\partial h_\tau}{\partial \tau} \big\vert _{\tau=\theta}$ |
| --- | --- | --- |
| Average | $\frac{N-1}{N}\theta + \frac{1}{N}\tau$ | $\frac{1}{N}$ |
| Median | $\theta \quad (\forall N \geq 3)$ | 0 |
| Random dictator | $\frac{N-1}{N}\theta + \frac{1}{N}\tau$ | $\frac{1}{N}$ |
| Novelty | $\tau$ | 1 |

We test these predictions in Fig. 3 by conducting simulations fixing *N* = 100 and varying the SCF. The simulations confirm the qualitative ranking implied by Table 1: *novelty* converges rapidly, *average* and *random dictator* improve slowly at nearly identical rates, and *median* shows little to no systematic improvement. Taken together, these results show that the same evolutionary process can produce radically different rates of improvement depending on the SCF.

**Figure 3.**
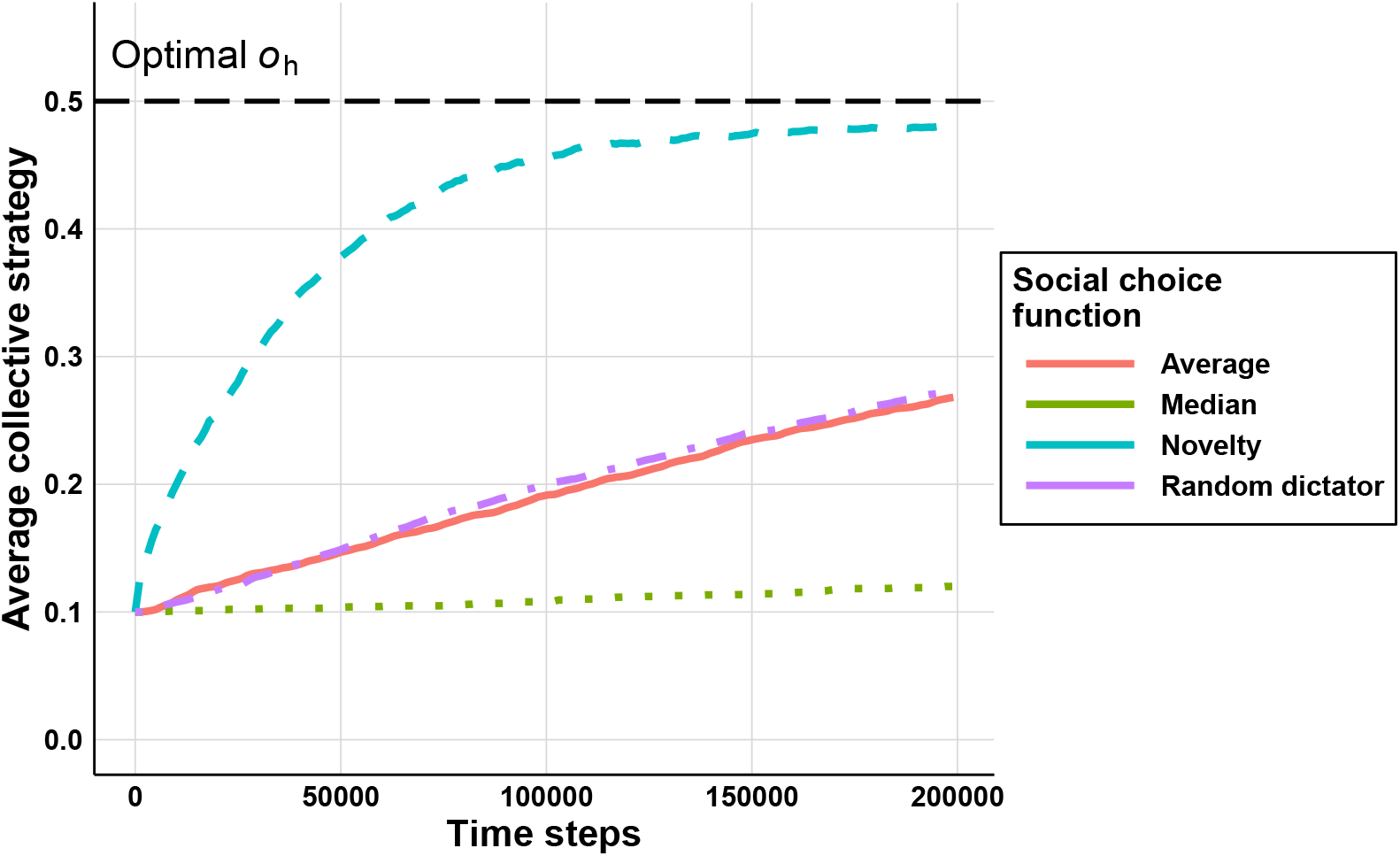
Evolution of mean collective strategy 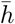 over time for four different social choice functions (mean across 50 replicates). We consider a population partitioned into *M* = 100 groups, each containing *N* = 100 individuals. Initial preferences are *x*(0) = 0.1. The payoff function has a single optimal value *o*_h_ = 0.5 and width *σ*_h_ = 0.25. The rate of random modifications is *µ*_m_ = 1 *×* 10^−5^, and the SD of random modifications is *σ*_m_ = 0.01.

#### Convergence to the same long-run outcomes

Finally, our results also predict that whenever marginal influence is positive, the population should converge to the same long-run equilibrium. This is difficult to assess in the simulations above, because they were designed to compare rates of change under rare, small random modifications, which makes convergence extremely slow. We therefore examine long-run outcomes separately in Appendix C, using larger and more frequent random modifications. These simulations confirm that populations converge towards the same equilibrium across group sizes and across the three SCFs with positive marginal influence, while preserving the earlier qualitative ranking in rates. The apparent exception is the *median* rule: under the small-variance approximation its marginal influence is zero, but once variation becomes larger it behaves more like an averaging rule, so it too converges towards the same equilibrium.

#### Additional checks

We examine whether these conclusions extend beyond the main simulation setting. We allow for preferential transmission within the same group in Appendix C.2, in which case we recover both the same qualitative ranking across SCFs and the same long-run convergence. We also turn to an explicit economic game, namely Hawk–Dove, where payoffs arise from an economic game rather than directly from an abstract payoff function. As shown in Case study 1 and Appendix E.1, the same long-run convergence still holds.

**Case study 1: Hawk–Dove and group conflict** The Hawk–Dove game is a standard game to model conflict within and between groups. In this game, pairs of players (individuals, or groups in our case) meet at random and compete over an indivisible resource. Each can play Hawk (conflict) or Dove (avoid conflict). A Hawk facing a Dove obtains the resource, but costly fights occur when both players choose Hawk. In the collective version, the players are groups, and the group strategy is determined by aggregating the preferences of group members (see Appendix E.1), which are then transmitted following the process described in Section 2.2.3. Our analysis predicts that, in the long run, political preferences and hence collective strategies should converge to the same probability of conflict as in the standard evolutionary Hawk–Dove game with individual players. Results in Figure S3 confirm this prediction. We find that (i) political preferences converge to, and remain at, the same equilibrium value under any of the SCFs considered earlier, and (ii) this equilibrium coincides with the mixed strategy that evolves in the standard Hawk–Dove game where individuals are the players (Figure S3).

### 4.2 When collective strategies fail to improve

So far, our results show that the improvement of collective strategies can vary sharply across decision procedures, ranging from rapid change to little or no systematic improvement in the simple setting considered above. We now ask whether, in richer settings, these same differences can have more dramatic consequences and cause improvement to fail altogether. To do so, we consider two illustrative cases in which cumulative improvement is notoriously harder to sustain: evolution with (i) a double-peaked payoff function and (ii) biased modifications.

#### 4.2.1 More complex payoff function

We first run simulations with a more complex payoff function with two locally optimal collective strategies (Appendix D.1 and illustrated in Figure S2). In this setting, because selection proceeds through gradual, incremental change, populations can sometimes not improve further once they have reached a locally optimal strategy, that is a strategy that works better than any slightly different ones (a local optimum), but still performs worse than some other, more distant strategy (the global optimum). In our illustrative example, this corresponds to a case in which societies perform well under a low-tax regime, which preserves private incentives, or a high-tax regime, which supports valuable public goods, while intermediate tax levels perform less well.

Our results show that whether this outcome happens or not varies strongly across SCFs. Populations using the average function, especially when groups are large, are more likely to remain stuck with inefficient collective strategies, whereas populations using functions such as novelty or dictator can reach the best collective strategy. This suggests that, with realistic payoff functions with multiple optima, how decisions are taken can have much more dramatic consequences than just slowing or speeding improvement. Instead, it may determine whether populations ever reach efficient collective strategies, potentially generating large and persistent payoff differences between populations.

#### 4.2.2 In the presence of a maladaptive bias

Second, we examine the case where modifications during transmission are, on average, more likely to move political preferences in a maladaptive direction (Appendix D.2). Continuing our illustrative example, this captures a situation in which individuals hold a bias against taxation, even when intermediate levels of taxation are beneficial. We examine such biases because, beyond being simple and plausible (Buss, 2015; Rubin, 2003), they underpin a seminal result in cultural evolution that whether selection can overcome maladaptive biases depends on population size (Henrich, 2004). Thus, this provides a clear benchmark for assessing how far predictions from previous work carry over to collective strategies. To enable a direct comparison, we vary the same parameter of total population size.

As in our main results, we begin with the limiting case in which each group contains a single individual (*N* = 1), so that collective strategies effectively behave like individual traits and we should recover the standard prediction from cultural evolution. Given our learning stage, this case is equivalent to a single well-mixed population of size *M*, so increasing population size simply means increasing the number of “groups”. Our results show that here, collective strategies change in the direction of the bias in small populations, whereas they converge to the optimal value in sufficiently large populations (Figure 4, top right panel). This reproduces the classic result that only sufficiently large populations allow selection to overcome biased transmission and improve traits (Henrich, 2004).

**Figure 4.**
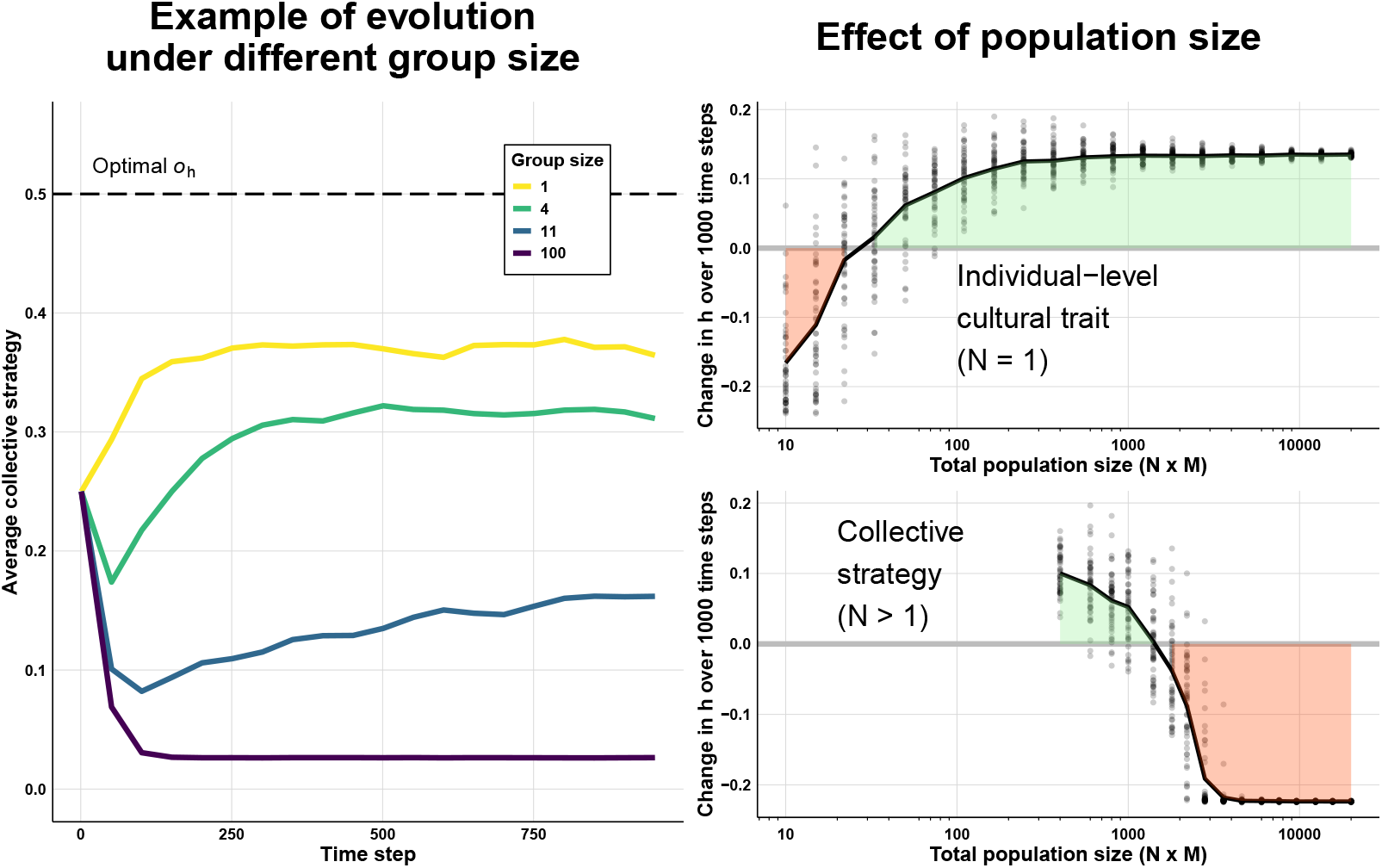
(Left) Evolution of the mean collective strategy 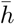 over time for group sizes: *N* = 1, 4, 11, 100. (Right) Change in the mean collective strategy after 1000 time steps, 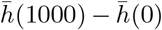, as a function of total population size, with initial values 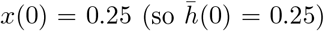. Dots are runs and lines are replicate means. The top row is the baseline *N* = 1, where population size is varied by changing the number of groups *M*. The bottom row is the collective case *N >* 1, where *M* = 200 is fixed and population size is varied via *N*. Random modifications occur at every transmission event (*µ*_m_ = 1) and the new value is drawn from a Gumbel distribution, truncated to [0, 1] with location (mode) equal to the previous value plus a bias δ_m_ = −0.01 and scale *σ*_m_ = 0.01. The payoff function has a single optimal value *o*_h_ = 0.5 and width *σ*_h_ = 0.25, so the bias is maladaptive at the initial condition.

We then turn to the case with a population structured in a fixed number of groups of size *N >* 1. Here the pattern is reversed. In small populations, collective strategies improve while in large populations, biased modifications accumulate and collective strategies converge towards a value which yields a low payoff (bottom right and left panel of Figure 4). This occurs because holding the number of groups fixed, increasing population size now corresponds to increasing group size, which as predicted weakens selection.

These results show that the way decisions are taken can affect not only the rate of change of collective strategies but also the long-run outcome. Case study 2 demonstrates the same point in a fully specified economic game by replicating and extending a model of evolution of institutions. More broadly, the results also show that the dynamics of collective strategies can differ sharply from those of models of a standard individual cultural trait.

**Case study 2: Evolution of a punishing institution** For our second case study, we revisit a classic model of evolution of institutions in which groups set a collective strategy to divide the benefits from a public good between punishing defectors and increasing group welfare (Powers and Lehmann, 2013). This model has been used to show how cooperation and group size can coevolve, enabling the emergence of large cooperative societies. We replicate these results (Appendix E.2), and, building on our insight that large groups experience weaker selection, we find that the collective strategy that evolves and persists at equilibrium in the original study is far from optimal. We further show that groups consistently adopt less efficient collective strategies as group size increases, while very efficient collective strategies persist if decisions are made using the novelty rule introduced earlier.

## 5 Discussion

A long-standing idea is that rules, institutions and other collective strategies may improve when political preferences associated with successful individuals are more often copied. This could generate efficient strategies even when individuals cannot evaluate their consequences, a phenomenon often seen as a key ingredient of human success (Henrich, 2016). Yet, whether this process reliably improves strategies, and under which conditions, remains unclear. We address this gap by analysing a general model of the evolution of political preferences and the strategies they determine. We summarise our findings in three points.

First, our results demonstrate that political preferences and collective strategies can indeed improve over time through selection, and across a wide range of conditions. In fact, our results show that for many ways in which groups could collectively decide on these collective strategies, and given enough time, collective strategies will converge toward the same locally optimal value. This process is sufficient for complex and efficient rules to emerge, even in the absence of individual foresight or causal understanding.

Second, our results also show that (i) the rate at which this improvement happens can vary dramatically, from populations that adapt quickly to others where the timescales required for improvement are implausibly long; and in some settings collective strategies may not improve at all, because selection is too weak to prevail against other forces such as maladaptive biases, or to dislodge populations from locally optimal strategies. While this usually happens because the link between success and transmission is weakened, for instance under a frequency-dependent learning bias (Boyd and Richerson, 1985; Henrich and McElreath, 2003; Aoki et al., 2011), here it is because the link between trait and success is weakened, as a person’s preference only partly shapes their group’s strategy.

Third, we identify the key factor that determines whether selection reliably improves collective strategies: the extent to which new preferences affect the strategies groups adopt. When this influence is diluted, for instance in large democratic groups, change can stall or drift; when it is stronger, selection drives rapid improvement in collective strategies. The intuition is that when individual influence is diluted, preferences for unfit collective strategies are more likely to persist because they are unlikely to be implemented. As we have shown, this marginal influence depends on many concrete features of groups, such as group size, political structure, or appeal to novelty, and suggests many other aspects of collective decision-making that could be explored.

Taken together, the key insight of our study is that collective strategies can improve over time through selection on political preferences, but whether they do so, and how effectively, depends critically on how collective decisions are made. This provides a qualified answer to a recent question in cultural evolution: whether institutions, often described as social technologies (Molho et al., 2024; Lie-Panis et al., 2024), can evolve through selection in ways analogous to material technologies. It also has a striking broader implication: because collective decision-making is itself often shaped by human-made rules, groups may partly determine the very selective forces acting on their own organisation.

These insights may also extend beyond humans, because the mechanism we analyse does not require individuals to understand or predict the consequences of alternative strategies. Many animal groups make collective decisions by signalling preferences that are then aggregated through informal rules (Conradt and Roper, 2005). Our results therefore suggest a simple prediction beyond humans: species or groups that differ in their decision making may differ in their capacity to produce efficient collective behaviours.

Our findings generalise the conclusions of previous models, which addressed important questions in other domains, by modelling the evolution of political preferences. These studies typically established their results under one or two collective decision procedures (Lehmann and Feldman, 2008; Powers and Lehmann, 2013; Johnstone et al., 2020; Mullon and Lehmann, 2022; Powers et al., 2023; Hunt et al., 2024), but we show that, given sufficient time, their conclusions extend across a much broader set of procedures. In fact, we show that even models treating groups as individuals, and ignoring how decisions are taken (e.g. Isakov and Rand, 2012; Gavrilets and Duwal Shrestha, 2021), yield the same evolutionary endpoints as if any procedure from this wide class were considered. An important methodological implication is that many questions about collective strategies that might seem to require complex, computationally heavy models can instead be studied using substantially simpler ones.

More generally, the idea that selection can improve collective arrangements has a long history, from early arguments about competition among firms and organisational forms (Alchian, 1950; Hannan and Freeman, 1984) to more recent work in cultural group selection (Boyd and Richerson, 2009; Smith, 2020). Our contribution is to clarify when such improvement should be expected once one makes explicit a plausible underlying mechanism, namely, that the strategies that successful individuals support are more often copied. To what extent this mechanism is implicitly assumed in these theories is not always clear. If it is indeed the case (for example, Smith (2020) cites models of this kind when illustrating cultural group selection), our results call for caution: selection may generate better collective strategies only under a narrower set of conditions than these arguments imply. If it is not, our results do at least underscore the importance of stating such pathways explicitly, echoing recent calls in the literature (Smaldino, 2014; Smith, 2020).

What does our model predict about change in collective strategies in real-world settings? At a minimum, our results identify when selection acting on political preferences could plausibly be an important driver of change, or when observed variation or change is more likely to reflect other processes. Put differently, it helps distinguish change driven by the differential transmission of political preferences from change driven by strategic adjustment or belief updating, processes that are often represented by different models, foreground different factors, and can generate different predictions. Before debating how much selection contributes in practice (Baumard and André, 2025), it is useful to establish when selection on political preferences can generate change in collective strategies at all.

At best, the model may offer a fairly direct account of how collective strategies change. This is most plausible in settings where people would revise their preferences less through careful deliberation than through copying others and trying things out. This may have been common in earlier societies, where records and formal knowledge were limited, and it can still occur today whenever collective strategies are too complex for their consequences to be understood. In such settings, the model shows how rules and institutions can improve over time even when individuals cannot reliably evaluate alternatives. It also identifies conditions that prevent such improvement, potentially explaining persistent disparities in success across populations or why some societies fail to adapt when new challenges make different rules more effective.

These cases also offer the best opportunities to test our predictions, either in historical settings or in the laboratory. The most informative natural cases are those that match the model’s group structure: multiple neighbouring groups facing similar problems over long periods, where success depends on developing effective organisation. Bronze Age Mesopotamian city-states, for example, experienced cycles of expansion and collapse often linked to failures to adapt institutions to environmental or demographic change (Frankopan, 2023). More recent ethnographic cases, such as the Turkana, organised into many semi-autonomous and competing groups, may provide this context (Mathew and Boyd, 2011). Similar conditions could also be created experimentally by letting multiple artificial groups make decisions and learn from one another’s performance, while ensuring that the consequences of their decisions remain hard to anticipate (as in experiments on other cultural traits; (Derex et al., 2019)).

Our analysis relies on two important simplifying assumptions. First, all groups in the population use the same social choice function, so a natural extension would be to allow it to differ across groups and potentially change over time. Second, strategies affect all individuals in a group equally. Relaxing this assumption is important, because many collective strategies benefit some individuals while disadvantaging others. But such cases would also require class-specific learning, since individuals disadvantaged by a collective strategy are unlikely to learn from those who benefit from it. We therefore leave this extension for future work. More generally, it would be valuable to explore learning rules beyond the payoff bias studied here, for instance the ones including biases identified in the literature (Henrich and McElreath, 2003), or learning directly from implemented collective strategies.

Taken together, our results show that the evolution of political preferences, and thus of the collective strategies they support, follows distinctive principles because collective strategies are actively constructed rather than directly inherited. This helps explain why insights from classical cultural evolution cannot be transferred uncritically, and instead points to the need for a wider research agenda on how political organisation, economic inequality, and the distribution of traits shape selection on political preferences and collective strategies.

## Code availability

All code necessary to reproduce the results is available at https://github.com/CedricPerret/InstDyn and archived at https://doi.org/10.5281/zenodo.21706203. Simulations were implemented using JuliassicPark.jl (Perret, 2026).

## Acknowledgements

I thank Thomas Currie for his feedback and for helping me develop the ideas presented here, and Laurent Lehmann for his advice on the analysis. I am grateful to Arthur Weyna for discussions on the analytical approach, and to Ludovic Maisonneuve for help with parts of the mathematics and for his comments on the manuscript. I also thank Miguel dos Santos, Simon T. Powers, and Charles Efferson for helpful comments and discussions.

## Appendix A Terminology

In this section, we briefly justify the choice of terminology used throughout this paper.

### Collective strategy

We use *collective strategy* to denote the strategy chosen at the group level, which determines or shapes group members’ actions in a subsequent economic game. We adopt this term to draw a parallel with the standard game-theoretic notion of an *individual strategy*, which is also typically the evolving trait in models of social evolution. We avoid the terms *institutions, rules*, or *regulations*. Although these are forms of collective strategies, they do not naturally cover cases where the collective strategy directly determines what group members do, such as the tactics of a sports team or the actions of an animal group. We also avoid *collective action*, which has an established meaning in the social sciences (Olson, 1965), and *norms*, which commonly refer to equilibria of games that emerge without an explicit collective decision (Young, 1998). That said, there might be overlap in the literature as a recent review on the emergence of norms (Smith, 2020) cites, as a study on the origin of norms, the exact same type of model that we extend and generalise in our paper.

### Political preferences

We use *political preferences* to denote individuals’ preferences over which collective strategy should be adopted. We use *preference* in the standard economic sense: a relatively stable ranking over alternatives. We qualify these preferences as *political* because they concern the actions of all group members, rather than only the actions of the individual herself. We avoid the term *opinion*, which in political science and opinion dynamics often refers to comparatively short-lived, context-dependent views (e.g. Castellano et al., 2009).

## Appendix B Complete evolutionary analysis

### Appendix B.1 General model

This section provides the full evolutionary analysis under the analytical approximation introduced in Section 2.3.1. We first treat the case where payoff may depend on the collective strategies of multiple groups (between-group interactions are allowed). We then recover the two special cases presented in the main text: (i) payoffs that depend only on the focal group’s collective strategy, and (ii) the baseline of groups of size *N* = 1.

#### Appendix B.1.1 Payoff function

In this section, we consider the payoff function

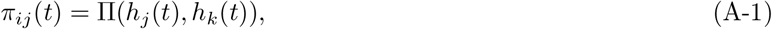

which denotes the payoff obtained by an individual *i* in group *j* when its group uses collective strategy *h*_*j*_(*t*) and all other groups use the same collective strategy *h*_*k*_(*t*). We will consider this payoff only in an approximation where all individuals other than the focal have the same political preference, and thus all groups other than the focal’s group adopt in expectation the same collective strategy. Thus, it is sufficient to write payoffs as a function of the pair (*h*_*j*_(*t*), *h*_*k*_(*t*)) rather than of the full vector of collective strategies of each group.

#### Appendix B.1.2 Small-variance approximation

##### Tracking political preferences to track collective strategies

To obtain analytical results, we adopt a standard small-variance approximation (Avila and Mullon, 2023). We assume that random modifications are rare and their effects are small, so that the distribution of political preferences is approximately Normal with mean *θ*(*t*) and small, roughly constant variance *V*_*x*_ (the variance being maintained by a balance between stabilising selection and a constant influx of variation).

Under this approximation, we treat the population mean collective strategy 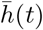 as being well approximated by the collective strategy associated with the average political preference, that is,

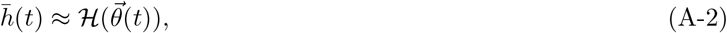

where 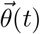 is the vector of size *N* where all entries equal the average political preference *θ*(*t*). By the unanimity property, 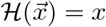 whenever all entries of 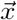 equal *x*, so this reduces to 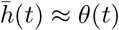. Therefore,

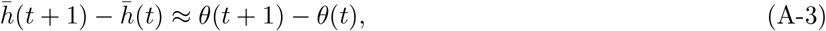

so tracking change in the average collective strategy reduces to tracking change in the average political preference.

##### Dynamics of political preferences

Under our assumptions, the change in average political preference obeys Mullon and Lehmann (2019)

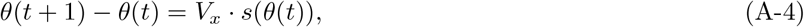

where *s*(*θ*(*t*)) is the selection gradient at time *t*. As the variance in political preferences is constant, computing the direction and amplitude of change in the average political preference reduces to computing the selection gradient. Denoting the political preference of a focal individual by *τ* and its fitness *w*(*τ, θ*), the selection gradient is

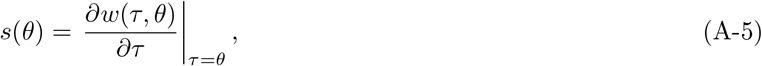

where for simplicity, we now omit the dependence on time (*t*) in this expression and in the following sections. The selection gradient describes the first-order marginal effect of an (infinitesimal) change in the political preference of the focal individual away from the population mean *θ*, on the fitness of this focal. The fitness of the focal is its expected number of copies over one time step.

#### Appendix B.1.3 Fitness of focal individual

The fitness of the focal individual is obtained by multiplying the probability that the focal preference is sampled in a given draw in eq. (4) by the total number of replacement draws, *MN*. This probability for the focal is obtained by substituting eqs. (1) and (A-1) into eq. (4) to express it as a function of political preferences, and then setting the preference of individual *i* to *τ* and the preferences of all other individuals to *θ*. This yields

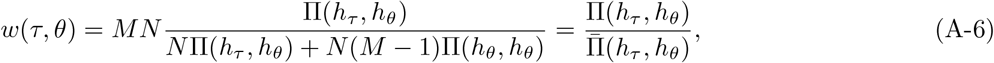

Where

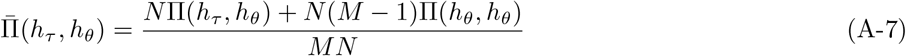

is the mean payoff in the population,

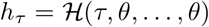

is the collective strategy in a group where one individual has preference *τ* and all others have preference *θ*, and

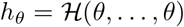

is the collective strategy in a group where all individuals have preference *θ*.

Assuming that the number of groups is sufficiently large that the focal group has a negligible effect on average population success, we can approximate 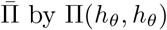, which yields

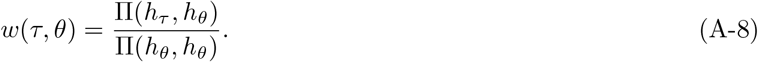

### Appendix B.2 Selection gradient

Using this expression for the fitness of focal individual, we can derive the selection gradient

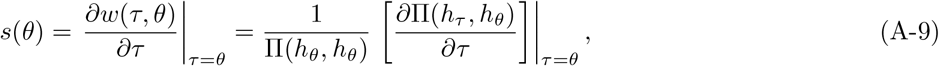

where we assume that the payoff function Π is twice continuously differentiable with respect to its arguments. Because *τ* affects payoff only through *h*_*τ*_, the selection gradient can be rewritten using the chain rule as

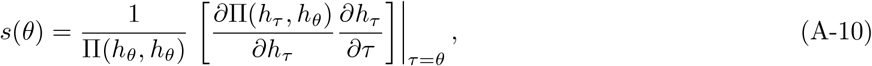

where 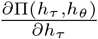 is the effect of a marginal change in the collective strategy on the focal individual’s payoff (the gradient with respect to collective strategies), and 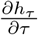 is the effect of a marginal change in the focal’s political preference on its group strategy (the *marginal influence*).

### Appendix B.3 Singular points

Singular points *θ*^∗^ are values at which the selection gradient vanishes. They are of particular interest because they are the candidate equilibrium outcomes of evolution, and their local properties determine whether evolution converges to them and remains there, moves away from them, or leads to evolutionary branching. Examining when the selection gradient in eq. (A-10) is equal to zero shows that singular points in the present model satisfy either

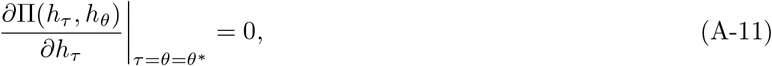

meaning that a marginal change in the average collective strategy would not change the payoff of the focal individual, or

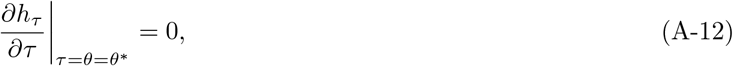

meaning that marginal influence is zero.

In the remainder of the paper, we focus on the first case and assume a positive marginal influence; singular points are therefore defined by a null gradient with respect to collective strategies. To determine whether the dynamics converge to these points, and how they behave once reached, we now turn to the stability analysis.

### Appendix B.4 Evolutionary stability

Evolutionary stability describes whether a population of individuals with political preferences at a singular point can resist invasion by individuals with slightly different preferences. Evolutionary stability is determined by the sign of the second derivative of the fitness of a focal individual with respect to its trait *τ* :

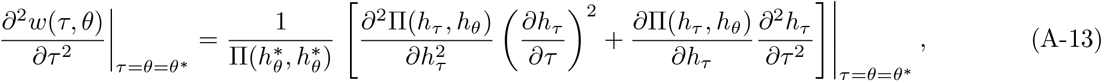

where 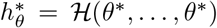 is the value of the collective strategy in a group composed exclusively of individuals with political preferences at the singular point *θ*^∗^. Recall from eq. (A-11) that a singular point is characterised by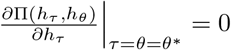. Substituting this condition into the previous equation yields

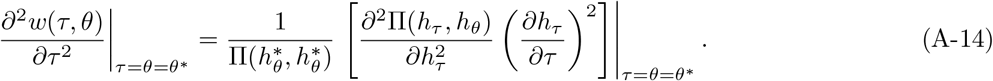

Since 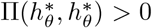, the sign of this expression is determined by the curvature of the payoff function with respect to the collective strategy,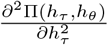, evaluated at *τ* = *θ* = *θ*^∗^.

A political preference is evolutionarily stable if this second derivative is negative. In other words, the induced collective strategy must locally maximise payoff for the focal group, holding fixed the collective strategy 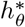 used by all other groups. This is the same condition for evolutionary stability as in a model where collective strategies evolve or groups reproduce directly as wholes.

### Appendix B.5 Convergence stability

Convergence stability describes whether a population that is initially away from a singular point will evolve towards or away from that singular point. The convergence stability is determined by the sign of the derivative of the selection gradient with respect to the average trait 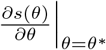. Using *s*(*θ*) described in eq. (A-9) and imposing the singular-point condition 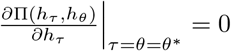 yields

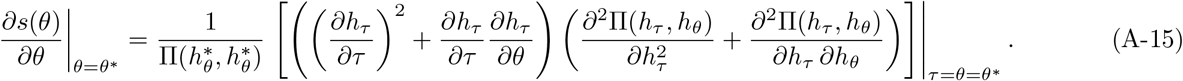

Under our assumptions of positive marginal influence and positive payoff, all terms outside the last parentheses are positive, so the sign of this expression is determined by this term. A singular point is convergence stable if

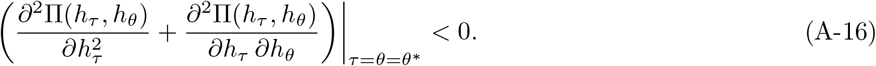

The first term, 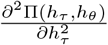, is the same derivative that determines the evolutionary stability. The cross-partial 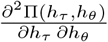 measures how the payoff of the focal individual changes when the collective strategy in the group containing the focal individual is increased while the collective strategy used by the other groups (i.e. the average collective strategy) is decreased, and *vice versa*. When this cross-partial is negative at the singular point, the payoff of the focal would increase if its group has a different collective strategy than other groups (negative frequency-dependent selection at the level of collective strategies).

### Appendix B.6 Classification of singular points

We can now classify singular points into the standard cases (Geritz et al., 1998), knowing that evolutionary stability is determined by the sign of the first term 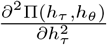 in eq. (A-16), while convergence stability is determined by the sign of the sum of this term and the cross derivative, 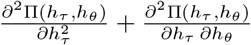,evaluated at the singular point.

First, if the singular point is not convergence stable (the sum of the two terms is positive), the point is a repellor and selection drives the population away from it for nearby trait values.

Second, if the singular point is convergence stable and evolutionarily stable (the first term is negative and the sum of the two terms is also negative), it is a continuously stable strategy (or attractor). Evolution drives the population towards this value of political preference (and thus of collective strategy) and keeps it there.

Third, if the singular point is convergence stable but not evolutionarily stable (the first term is positive, while the sum of the two terms is negative), it is an evolutionary branching point. The population is first attracted to this value of political preference, but once close to it, selection becomes disruptive and leads to the emergence of two distinct, stable political preferences. Depending on how political preferences are aggregated into a collective strategy, this may in turn generate two or more than two collective strategies. Branching happens only if the cross-partial derivative is sufficiently negative, meaning that there is a strong payoff advantage for a group to adopt a collective strategy that differs from the collective strategies used by other groups.

Finally, in the absence of a singular point, selection pushes political preferences, and hence collective strategies, consistently in one direction either upward or downward (depending on the sign of the selection gradient) until they reach a boundary.

### Appendix B.7 Special cases

#### Appendix B.7.1 In the absence of group interactions

In the main text we restrict attention to the case in which there are no group interactions and thus payoffs depend only on the collective strategy implemented in the focal group. Under this assumption, the payoff function can be written as

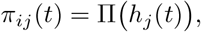

with no explicit dependence on the collective strategies used in other groups. Because the selection gradient and evolutionary stability depend only on derivatives of the focal individual’s fitness with respect to its own trait, dropping the second argument of Π leaves the expressions for the selection gradient, singular points, and evolutionary stability unchanged, except that Π(*·, ·*) is replaced by Π(*·*).

#### Appendix B.7.2 Baseline case with groups of 1

In the main text, we consider a baseline case where group size *N* = 1 and thus, each individual constitutes its own group. More generally, this baseline represents any case in which the evolving trait maps directly to the implemented collective strategy and hence to fitness. This includes both the standard case of an individual trait and the case in which transmission is assumed to be between groups (or groups themselves reproduce), so that the transmitted and evolving trait is directly the collective strategy. In this case, the implemented collective strategy of the focal individual equals its political preference, *h*(*τ, θ*) = *τ*. The marginal influence is therefore

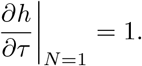

Substituting this into the general expression for the selection gradient (eq. (A-10)) yields the expression presented in Section 3 of the main text. The conditions for the singular point, evolutionary and convergence stability do not depend on the values of the marginal influence, or on *N* more generally, and thus remain the same as in previous sections.

## Appendix C Additional simulations and robustness checks

In this section, we extend the simulations presented in the main text in two ways. First, we consider larger and more frequent random modifications. Second, we allow for transmission to preferentially happen within groups.

### Appendix C.1 Larger and more frequent random modifications

In the main results, random modifications occurring during transmission of political preferences are assumed to be rare and small, which allows us to run simulations in conditions close to the approximation made to obtain analytical results. However, given that political preferences are cultural traits, random modifications in real populations may occur more frequently and with larger effect. We therefore rerun the same simulations, both for populations using the average as the SCF (with groups of different sizes) and for populations using the alternative SCFs considered in the main text, under a hundredfold higher rate of random modifications, *µ*_m_ = 0.001, and a larger SD of random modifications, *σ*_m_ = 0.1. These results reproduce Figures 2–3; they are shown in the left panel of Figure S1 and discussed in Section 4.1 of the main text.

### Appendix C.2 Allowing learning to occur preferentially within groups

In the main model, the probability that a political preference is sampled to replace an existing preference does not depend on whether the individuals carrying them belong to the same group or to different groups. We now relax this assumption and consider a setting where transmission may occur preferentially within or across groups, controlled by a parameter *m*.

#### Appendix C.2.1 Model definition

Formally, we retain the life cycle described in Section 2.2, but modify the learning stage (iii) as follows. The probability that individual *i* of group *j* is copied by an individual of group *l* is

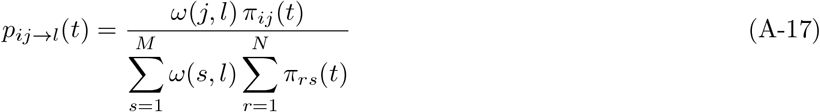

Where

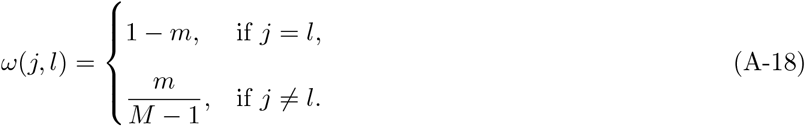

Thus, individuals from the same group and from other groups compete in a single sampling process, but their payoffs are weighted according to their group of origin. The parameter *m* controls the relative contribution of individuals from outside the focal group, rather than directly fixing the proportion of replacements originating from other groups.

#### Appendix C.2.2 Interpretations

This model admits two interpretations, depending on whether the transmission step is considered to represent vertical learning or horizontal learning.

##### Vertical learning

Under a reading in which individuals reproduce and offspring learn from their parents, the model corresponds to a life cycle in which offspring disperse after reproduction and then all offspring (philopatric and migrants) compete for positions within groups. Here, *m* is a literal migration rate: the probability that an offspring disperses to a different patch. The model in the main text corresponds to the well-mixed case (*m* = 1 − 1/*M*) of the present model, a standard baseline in evolutionary theory (Rousset, 2004).

##### Horizontal learning

Under a reading in which individuals remain in place and update their political preferences through horizontal social learning, the model corresponds to a life cycle where learners select a model based on their payoff and on whether the model is from the same group. In this case, *m* is the relative attention learners give to models outside their own group: values below 1 − 1/*M* correspond to a bias towards copying group members, and values above it to a bias towards copying outsiders. The model in the main text corresponds to the case with no group bias: individuals choose whom to learn from solely on the basis of payoff, neither favouring nor avoiding members of the same group.

#### Appendix C.2.3 Analysis

We run simulations using a low value of parameter *m* = 0.2. This either corresponds to limited dispersal between groups or more frequent within-group learning, justified by more frequent interaction or exchange of information within groups. We assess this case under the same higher rate of random modifications and higher SD of random modifications regime as above. The reason is that, when transmission occurs mainly within groups, most learning events involve individuals who experience the same collective strategy and hence have the same payoffs. Under rare, small random modifications, this makes the dynamics largely dominated by quasi-neutral resampling, so differences between SCFs are difficult to detect.

#### Appendix C.2.4 Results

The outcomes are presented in the right panel of Figure S1 and discussed in the last paragraph of Section 4.1 of the main text.

**Figure S1.**
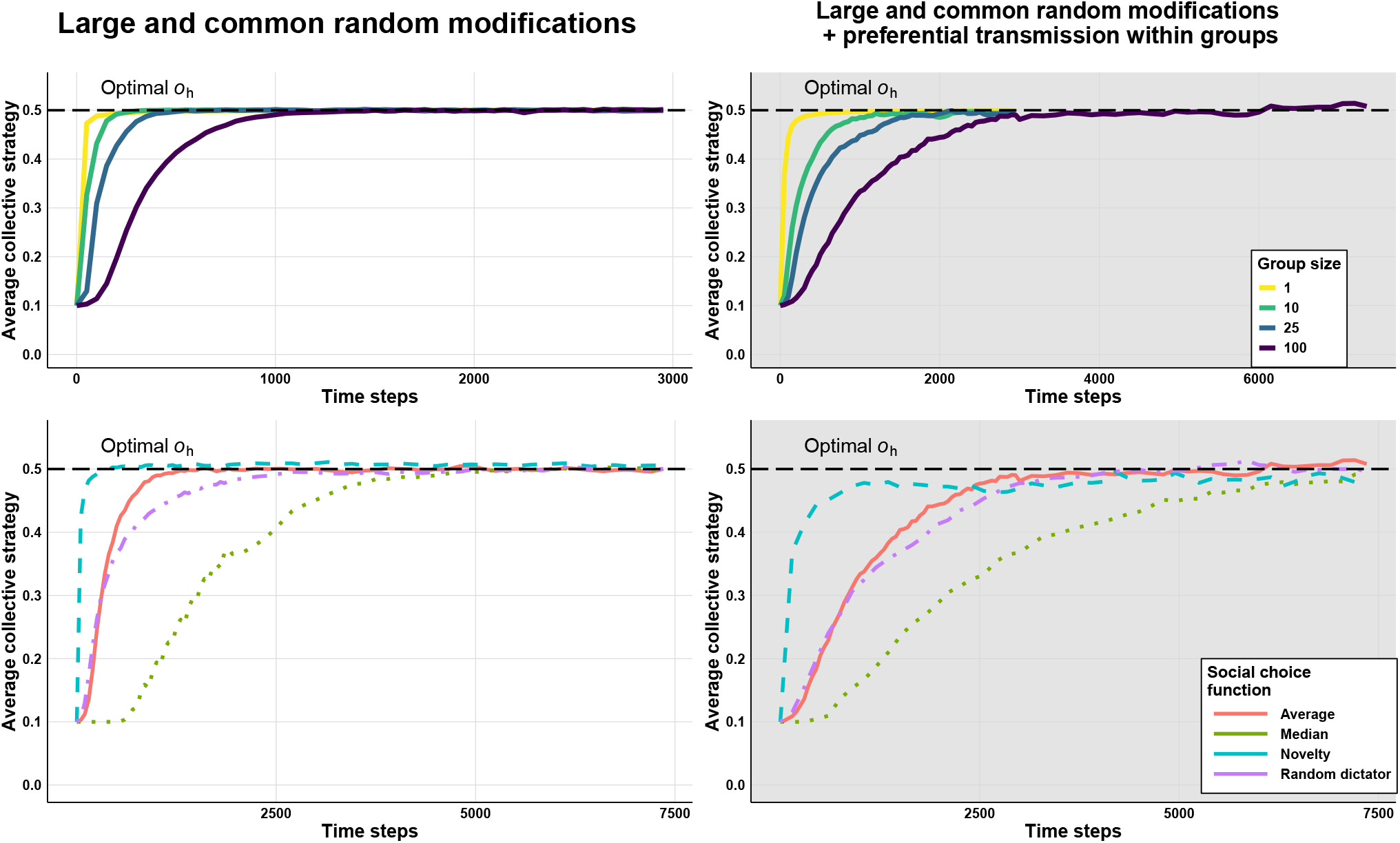
Evolution of collective strategies *h* over time, averaged across 50 replicates, for different group sizes when the social choice function is the average of political preferences (top), and for different social choice functions for group size *N* = 100 (bottom). The left panel shows simulations with a higher rate of random modifications (*µ*_m_ = 0.001) and larger SD of random modifications (*σ*_m_ = 0.1); the right panel shows simulations with the same parameters combined with preferential learning within groups (*m* = 0.2). We consider a large population of fixed size (10 000), partitioned into groups of size *N*, so that the number of groups is *M* = 10 000/*N*. Initial preferences are *x*(0) = 0.1. The payoff function has a single optimal value *o*_h_ = 0.5 and width *σ*_h_ = 0.25.

## Appendix A Simulations in richer settings

### Appendix D.1 Payoff functions with multiple optima

#### Appendix D.1.1 Introduction

So far, we assumed a payoff function characterised by a unique collective strategy providing maximum payoff. In these cases, the fact that strategies change in the direction that increases payoff locally (i.e. where a small change in that direction would increase payoff) implies that the strategies will eventually reach this optimal value. If the relationship between collective strategy and payoff is more complex, local improvement over time does not guarantee that. Populations may instead become stuck with a strategy that works better than any slightly different ones (a local optimum), but still performs worse than some other, more distant strategy (the global optimum). We therefore conduct simulations assuming a more complex payoff function with two optimal values.

#### Appendix D.1.2 Method

In this section, we consider a payoff function defined as the sum of two Gaussian functions:

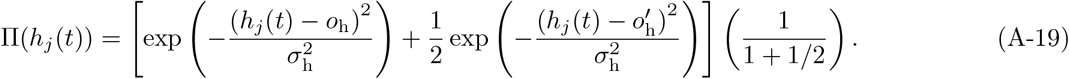

The first Gaussian function, centred at *o*_h_, defines the global optimum, that is the collective strategy that provides the maximum payoff. The second Gaussian function, centred at 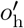, represents a local optimum, that is the collective strategy that yields a higher payoff than any nearby strategies, even though it is not the best overall. We assume that the global optimum provides a payoff twice higher than that of the local optimum. Returning to the taxation example, this type of payoff function could be justified on the basis of empirical evidence and theoretical work suggesting a more complex relationship between tax rates and public revenue than the single-peaked function considered earlier. For instance, Tavor et al. (2022) show that under certain economic assumptions, the Laffer curve can exhibit two distinct optima, as considered here.

#### Appendix D.1.3 Simulation procedure

We know from our previous results that, given sufficient time, the population will eventually reach one of the optima. A more informative way to assess the capacity of collective strategies to improve in such a case is to quantify the population’s ability to escape the local optimum and reach the global optimum. To do so, we conduct simulations in which all initial political preferences are initialised at the local optimum 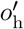 and record, across 100 replicates, the proportion of populations that escape the local optimum within 1,000 time steps.

We repeat this analysis for local optima that differ in how difficult they are to escape, which we refer to as the strength of *lock-in*. To vary lock-in (see top panel of Figure S2), we position the two optima so that the distance from the local optimum to the nearest point of equal payoff is controlled by a parameter *d >* 0

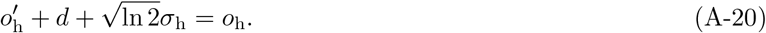

Beyond this point, selection will ensure that the collective strategy moves uphill toward the global optimum. This distance is sometimes called the payoff valley and *d* measures the distance to cross this valley.

We run simulations for a range of values of *d*. We choose parameters on the random modifications (*µ*_m_ and *σ*_m_) such that populations in the baseline (*N* = 1) can escape strongly locked-in optima. We then examine populations with different group sizes or using different social choice functions.

**Figure S2.**
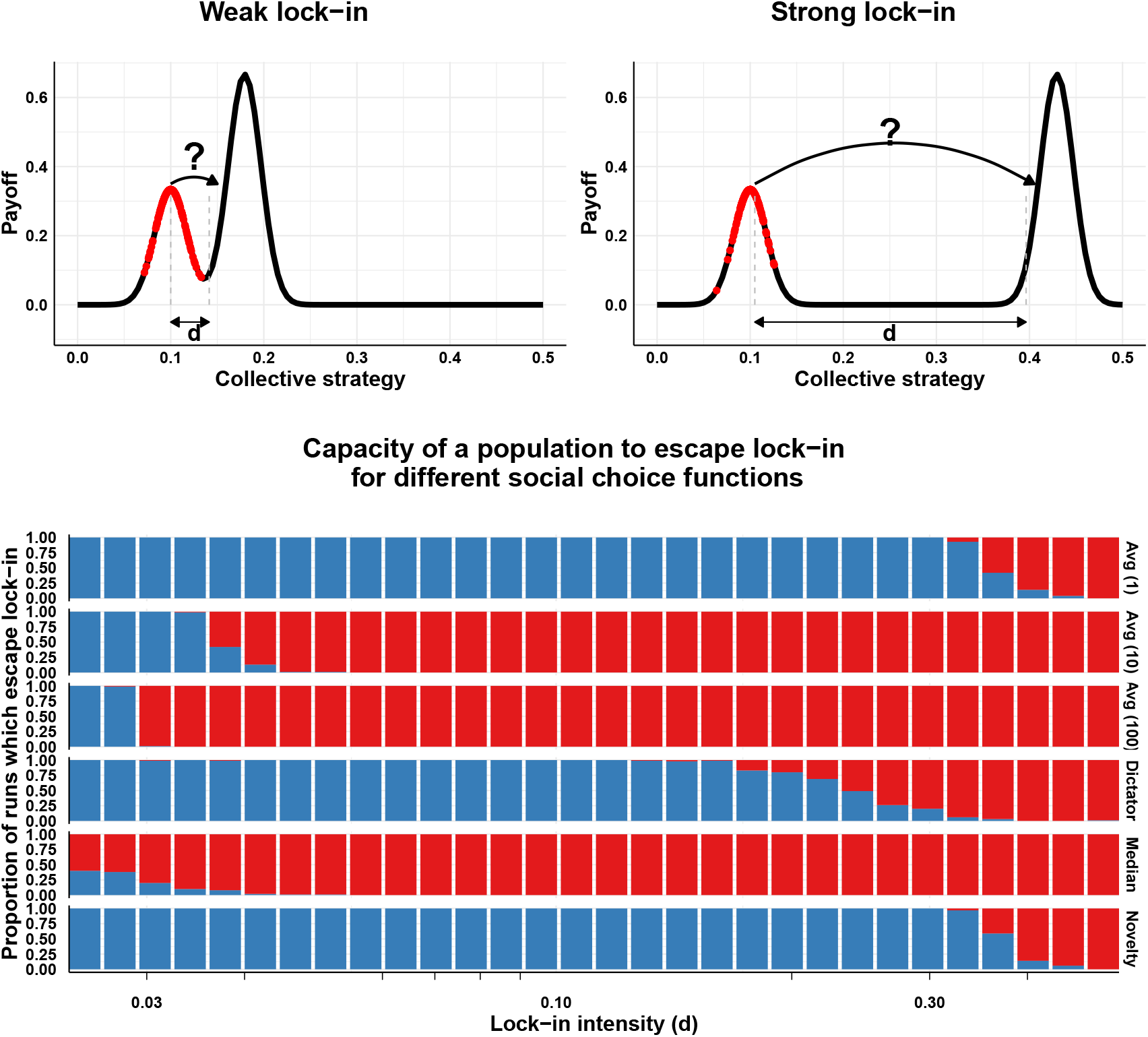
Ability of populations to escape a locally optimal collective strategy for different social choice functions or group sizes. Escape success is measured as the proportion of replicate runs (100 replicates) in which the population escapes the initial local optimum within 1,000 time steps, as a function of the distance *d* between the two optima. A higher distance represents a stronger lock-in, making transitions more difficult. The top panel illustrates the setup. Populations are initialised at the lower local optimum, located at 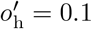. The question is whether the population (red) can cross the valley and reach the higher optimum. A run is counted as having escaped if the population mean collective strategy exceeds 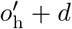. The bottom panel shows the proportion of runs that escape, plotted against *d*. Unless otherwise specified, parameters are *σ*_h_ = 0.025, *M* = 10 000/*N, N* = 100, *µ*_m_ = 0.001, and *σ*_m_ = 0.1.

#### Appendix D.1.4 Results

##### Evolution for different group sizes under averaging

We first consider populations in which the social choice function is the average of political preferences, while varying group size. Increasing group size (first three rows of Figure S2) greatly reduces the population’s ability to escape the local optimum. Thus, besides slowing the improvement of collective strategies, larger groups also make populations more likely to remain trapped at locally optimal collective strategies. This is because, in larger groups, a change in a single individual’s political preference produces only a small change in the group’s collective strategy. In the limit of very large groups, the influence of any one individual becomes negligible. Additionally, as more individuals simultaneously change their political preferences, these shifts tend to cancel out on average, further limiting the effect of individual change on the collective strategy.

##### Evolution for different social choice functions at fixed group size

Different SCFs also strongly influence the ability of populations to escape local optima (Figure S2). The novelty rule performs best, allowing populations to reach distant optima about as reliably as in the baseline case of groups with a single individual. It is followed by the dictator rule, then the median rule, both of which outperform the average rule in large groups. Notably, the dictator rule performs better here, even though it previously performed about as well as the average rule in terms of the rate of change. This is because most changes in political preference do not affect the collective strategy, but when the dictator changes their preference, the group can immediately adopt a very different strategy. This allows larger jumps and increases the chance of escaping local optima. These findings show that outcomes on complex payoff functions are not always intuitive, and that results from simpler settings do not always carry over. They also point to a broader dimension of evolvability: not only the rate of incremental improvement matters, but also the ability to make larger jumps and escape local traps. Understanding how different SCFs balance these two dimensions remains an important question for future work.

### Appendix D.2 Evolution in the presence of biased modifications

This section describes the implementation of biased modification used in Section 4.2.2. We follow the standard formulation introduced by Henrich (2004), adapting it to our framework.

#### Appendix D.2.1 Model definition

We implement the same life cycle as in our main simulations, with one difference: when a random modification occurs, the updated value of political preference is drawn from a Gumbel distribution, truncated to [0, 1], with location (mode) equal to the previous value plus a bias δ_m_ *<* 0 and scale *σ*_m_. Because populations are initialised below the optimum (*h*_*j*_(0) = 0.25 for *o*_h_ = 0.5), this makes modifications maladaptive on average. Furthermore, to replicate the modelling choices of Henrich (2004), we assume that modifications occur at every transmission event (*µ*_m_ = 1).

In the baseline case, we set *N* = 1, so each group contains a single individual and collective strategies effectively behave like individual traits: each political preference maps directly to payoffs. With our learning stage, this is equivalent to a single well-mixed population of size *M*, so increasing population size simply corresponds to increasing *M*.

In the extension with *N >* 1, individuals are organised into groups and collective strategies are collectively determined within each group. In this case, an increase in total population can reflect either more groups or larger groups. Here we fix the number of groups, which can be interpreted as a fixed number of sites (e.g. territories or resource patches), so larger population size corresponds to increasing group size *N*.

#### Appendix D.2.2 Results

The results are described and discussed in Section 4.2.2 in the main text.

## Appendix E Case studies

In this section, we provide the detailed model specification and results for the case studies used in the main text. We begin with an extension of the Hawk–Dove game, a canonical model of conflict (Maynard Smith and Price, 1973), before turning to a more complex model on the evolution of institutional rules (Powers and Lehmann, 2013).

### Appendix E.1 Hawk–Dove game

#### Appendix E.1.1 Our implementation

We assume a population subdivided into groups. At each time step, every group initiates *k* encounters with other groups chosen at random (with replacement). During each encounter, the two groups play a classic Hawk–Dove game, in which each chooses between two strategies: Hawk (escalate conflict) or Dove (avoid escalation). All individuals within a group receive the same payoff, determined by the group’s strategy and that of its opponent according to the payoff matrix

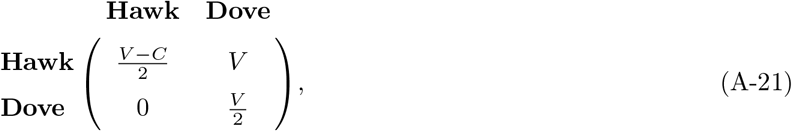

where the entry in row *X*, column *Y* gives the payoff of a player using strategy *X* against an opponent using strategy *Y*. To avoid negative fitness values, all payoffs are shifted by adding a constant of 1. Payoffs are then normalised by the number of encounters, since groups may not all participate in the same number of encounters.

We embed the Hawk–Dove game into the same simulation framework used in the main text. Each individual *i* in group *j* holds a political preference *x*_*ij*_ ∈ [0, 1], interpreted as their tendency to favour pacifist interactions. A group’s collective strategy describes the probability to play dove when meeting another group. It is generated by aggregating members’ political preferences using one of the social choice functions described in Section 4.1. Political preferences are transmitted following the same process as in the main simulations. This setup matches the formulation for instance used by Johnstone et al. (2020), who assume the equivalent of the random dictator rule.

#### Appendix E.1.2 Analysis

Our theoretical results in Section 3 make two predictions. First, for any SCF with positive marginal influence, selection drives political preferences and collective strategies to converge to the same long-run value. Second, this common endpoint should match the evolutionarily stable strategy of the individual-level Hawk–Dove game, namely a probability to play dove equal to 1 − *V*/*C* (equal to 0.5 under our parameters *V* = 1 and *C* = 2). To test these predictions, we initialise the population with political preferences far from the predicted equilibrium value (*x*_*ij*_(*t*) = 0.25) and track the evolution of the collective strategy for different SCFs. For reference, we also simulate the standard individual-level case by setting *N* = 1, where each individual’s political preference is their strategy.

**Figure S3.**
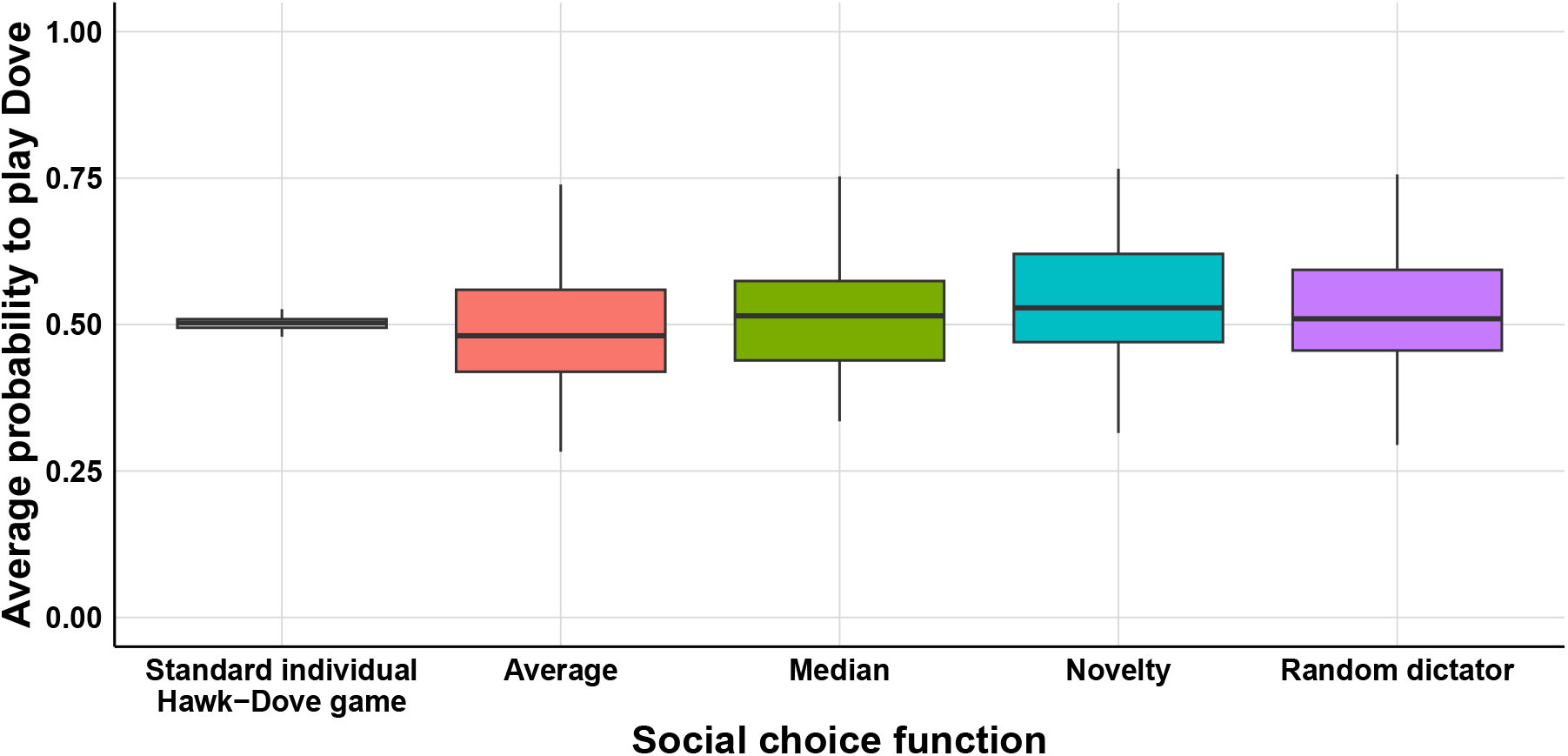
Average collective strategies at equilibrium under different social choice functions in a Hawk–Dove game. We compare four social choice functions with the standard individual-level Hawk–Dove game, recovered by simulating *M* = 10,000 groups of size *N* = 1, so that each individual’s political preference directly represents their strategy. In all other cases, the population is structured into *M* = 100 groups of size *N* = 100. Values are shown after 20,000 time steps and averaged across 50 replicates. Initial preferences are *x*(0) = 0.25. We set *µ*_m_ = 0.001 and *σ*_m_ = 0.1 so that populations approach equilibrium within computationally feasible times even when improvement is very slow. Each group initiates *k* = 10 encounters per time step, although groups may participate in more encounters overall when chosen by others. Game parameters are *V* = 1 and *C* = 2.

#### Appendix E.1.3 Results

Our results confirm that, although the rate of evolutionary change varies across SCFs, the long-term equilibrium value of the collective strategy is the same for all functions considered here (Figure S3). Moreover, this common endpoint matches the equilibrium predicted by the standard individual-level Hawk–Dove game. This holds despite the fact that groups, rather than individuals, are the strategic players in the interaction, and despite substantial differences in aggregation mechanisms. The results are further described and discussed in Case study 1 in the main text.

### Appendix E.2 The evolution of punishing institutions

We now replicate and extend a model of evolution of institutional rules developed by Powers and Lehmann (2013).

#### Appendix E.2.1 Summary of previous model

The model of Powers and Lehmann (2013) simulates a population structured into patches, each of which may contain social and asocial individuals. Social individuals form groups and engage in a public goods game, while asocial individuals do not. Social individuals may contribute to the public good or defect, in which case they get the benefits of the public good without paying a cost. Each group also adopts a collective strategy which sets the proportion of the public good invested in increasing the group’s carrying capacity while the rest is used to sanction defectors. Following the original terminology, we refer to collective strategies here and thereafter as *institutions*.

Each individual carries a trait setting their behaviour in social interactions, either asocial, cooperator, or defector, as well as a political preference over the institution. Individuals’ fitness depends on their type (cooperators and defectors pay a cost for implementing the institution and cooperators pay an additional cost to contribute to the public good), and carrying capacity (which depends on the public good) as per eq. (1) of the original paper. All individuals reproduce in proportion to fitness, and offspring inherit both trait and political preference, with rare small random modifications.

#### Appendix E.2.2 Our implementation

We reimplement the model as described in the original paper, using the same parameter values reported in Table 1 of Powers and Lehmann (2013), except for the treatment of mutations. Rather than drawing mutation effects from a normal distribution and clamping values at the boundaries (as in the original simulations), we implement mutations as draws from a truncated normal distribution. This avoids artefactual boundary effects, whereby mutation disproportionately pulls trait values toward the extremes, especially when variance is high. Truncated normal mutations ensure more realistic trait evolution and preserve the intended behaviour of the model. Second, we use a smaller migration rate (0.05), since cooperation fails to evolve under this mutation regime when the rate is 0.1. Importantly, the key qualitative result of the original paper—the coevolution of cooperation, institutions, and group size—still holds under this regime. Third, Powers and Lehmann (2013) assume that an offspring mutates with probability *µ*_m_ = 0.01, in which case one of its two traits, chosen at random, is modified. We instead mutate each trait independently with probability *µ*_m_ = 0.005, which gives the same expected number of mutations per trait. The two schemes differ only in that ours occasionally modifies both traits in the same offspring, which happens rarely enough to be inconsequential here. For all other aspects of the model, payoff calculation, demographic update, and life cycle, we follow the original specification.

Unlike in previous sections, comparing the speed of convergence toward the optimal institution is of limited interest, because initial conditions assume groups begin very small and consist entirely of asocial individuals. Under these conditions, political preferences drift randomly in the early generations and converge rapidly once social individuals emerge. We therefore focus on the long-run equilibrium values of the institution *h*.

#### Appendix E.2.3 Results

##### Typical dynamics

We begin by describing the typical dynamics reported in the original paper. Starting from a population of asocial individuals, cooperators can invade because they generate additional benefits through the public good, provided the institution in their groups allocates enough resources to punishment. This prevents defectors from obtaining a higher fitness than cooperators. Second, once cooperators are established, the value of the institution evolves to minimise waste. Punishment only needs to reduce defectors’ fitness below that of cooperators for cooperators to resist invasion from defectors. Any additional resources allocated to punishment have no effect and instead divert resources away from group growth. Thus, when defectors are rare, allocating too much of the public good to punishment is inefficient, since those resources could otherwise support group growth; but when defectors are common, allocating too little allows them to thrive. As a result, and because defectors are rare at equilibrium, selection tends to shift institutions toward allocating a large share of the public good to sustaining group growth. We start by reproducing these results in Figure S4. We recover the result that over time, asocials are replaced by cooperators, while these cooperators manage to avoid invasion by defectors. At the same time, groups grow to a large size, meaning that they are able to produce the public good steadily and suppress defection. The value of the institution also converges to a non-random value around 0.75, indicating that selection is acting. As explained in the original paper, this can lead to larger and larger societies, with the equilibrium group size depending on the potential maximum benefit provided by the public good (bottom panel).

**Figure S4.**
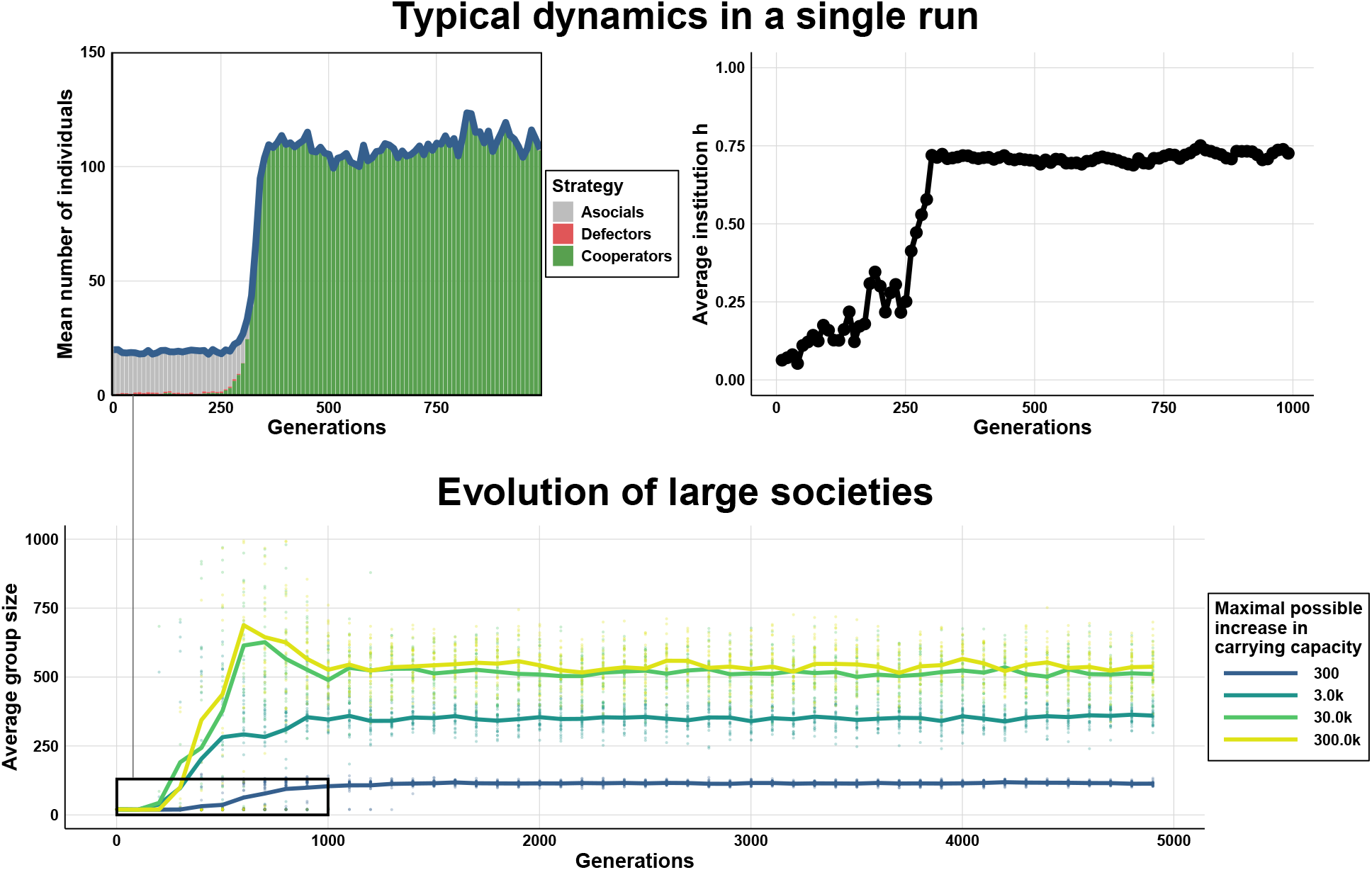
Replication of the model from Powers and Lehmann (2013). Top panels present the evolution of the average group size, group composition and institution over time for a single run (*β* = 300, *γ* = 7.5 *×* 10^−3^). Bottom panel shows the evolution of average group size over time for different equilibrium group sizes. As in the original paper (page 1360), we manipulate equilibrium group sizes by varying the demographic parameters *β* and *γ*, following the same scaling procedure (i.e. increasing *β* tenfold and decreasing *γ* tenfold). We reproduce the two cases from the original study (*β* = 300, *γ* = 7.5 *×* 10^−3^; *β* = 3,000, *γ* = 7.5 *×* 10^−4^), and we generate two additional cases (*β* = 30,000, *γ* = 7.5 *×* 10^−5^ and *β* = 300,000, *γ* = 7.5 *×* 10^−6^). All parameters match those used in the original study, except for the mutation regime (as described in the text): cost of cooperation *C* = 0.1, cost of institution *I* = 0.1, individual benefit from public good *B* = 0.9, base growth rate *r*_a_ = 2, asocial carrying capacity *K*_A_ = 20, per capita effect of social individuals on asocials *α*_sa_ = 0.05 and of asocials on socials *α*_as_ = 0.05, mutation rate *µ* = 0.005, mutation variance *σ*^2^ = 0.1, number of patches *N*_p_ = 50, and migration rate *m* = 0.05.

**Figure S5.**
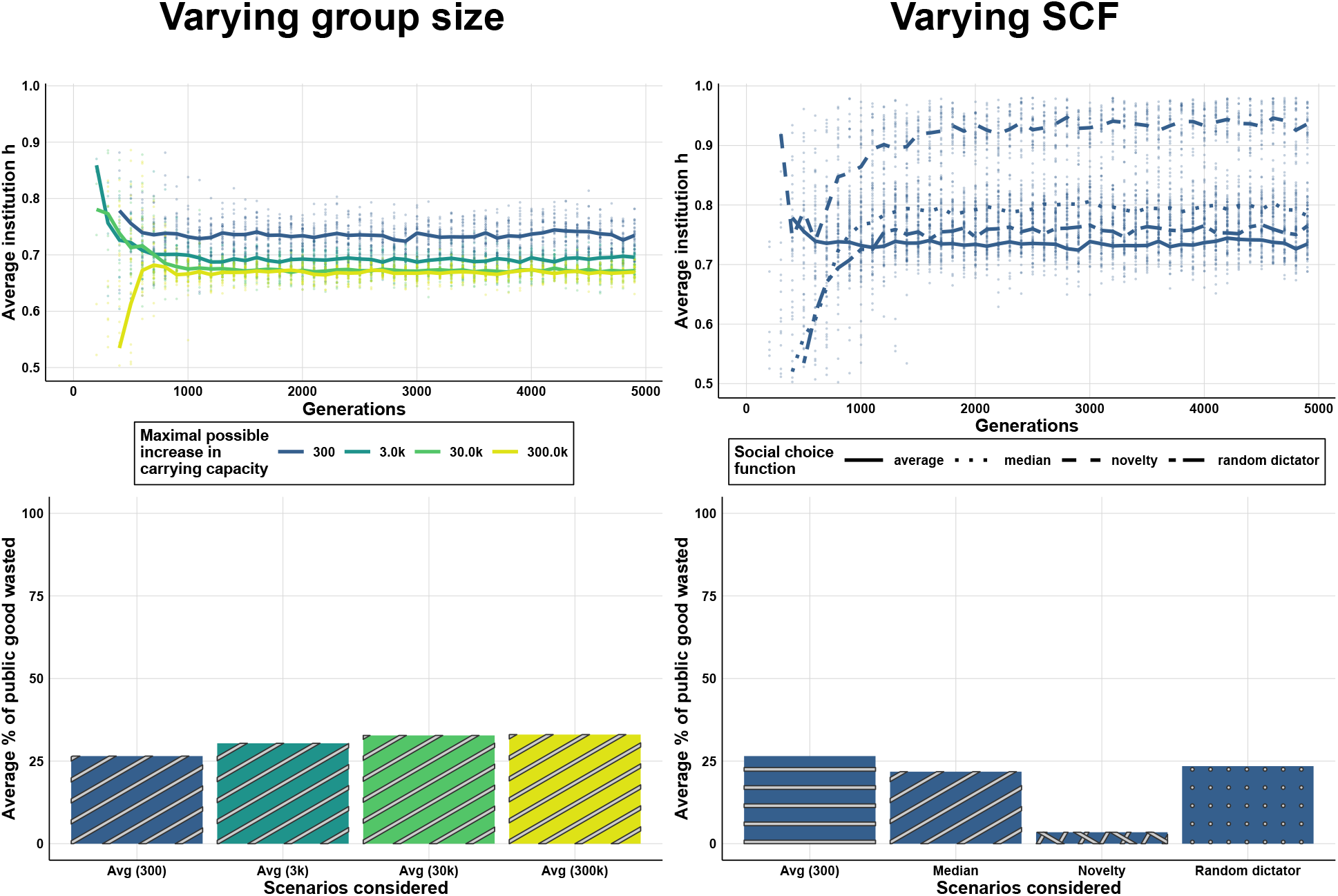
Evolution over generations of the institution *h* (top) and the fraction of the public good wasted (bottom) for different equilibrium group sizes and social choice functions. Points represent individual replicates; lines show the average across 30 replicates. The fraction of the public good wasted is calculated as 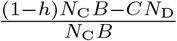 where *N*_C_ and *N*_D_ are the number of cooperators and defectors, respectively. The numerator is the amount invested in punishment minus the minimum amount needed to make defectors less fit than cooperators, and the denominator normalises this quantity by total public-good production. Positive values therefore indicate overinvestment in punishment. As in the original paper, we manipulate equilibrium group size by varying the demographic parameters *β* and *γ*, following the same scaling procedure (e.g. increasing *β* tenfold and decreasing *γ* tenfold) to generate two additional cases: *β* = 30,000, *γ* = 7.5 *×* 10^−5^ and *β* = 300,000, *γ* = 7.5 *×* 10^−6^. All parameters match those used in the original study, except for the mutation regime (as described in the text): cost of cooperation *C* = 0.1, cost of institution *I* = 0.1, individual benefit from public good *B* = 0.9, base growth rate *r*_a_ = 2, asocial carrying capacity *K*_A_ = 20, per capita effect of social individuals on asocials *α*_sa_ = 0.05 and of asocials on socials *α*_as_ = 0.05, mutation rate *µ* = 0.005, mutation variance *σ*^2^ = 0.1, number of patches *N*_p_ = 50, and migration rate *m* = 0.05.

##### Does selection fully optimise the institution?

Previously, there was no particular reason to question whether this value of institution was the most optimal one, as it showed signs of being selected and of becoming more efficient (reflected by growing group size). However, our earlier findings show that selection may be limited as groups are large and decisions are made by averaging. In particular, this study assumed mutations with large effects and strong drift (because the number of groups is small), which tends to pull the average value of the institution toward the centre of the trait space. This suggests that the value of the institution may, in fact, lie far from the optimum.

To test this, we measure the efficiency of the institution by the fraction of public good wasted, that is, the share of the public good devoted to punishing defectors above what is necessary to make them have a lower payoff than cooperators (see caption of Figure S5 for the formula). If selection operates effectively, institutions at equilibrium should allocate just enough to punishment to stabilise cooperation, while using the rest to increase the carrying capacity. In practice, however, the bottom left panel of Figure S5 shows that around a quarter of the public good is wasted. This indicates that the institution at equilibrium is far from optimal, consistent with our predictions. Selection is acting but it is not strong enough to counterbalance mutation and drift effects in this context.

##### Group size limits the evolution of efficient institutions

We then conduct a similar analysis to that used throughout the paper by examining the effect of varying group sizes (equilibrium group sizes in this case). Left panels of Figure S5 show that larger groups tend to evolve values of *h* closer to 0.5, and waste a larger fraction of the public good. These results are consistent with our analytical predictions: in large groups, the strength of selection is weak such that recurrent mutation maintains a broad distribution of preferences rather than one clustered around the favoured value. This, on average, yields institutions close to the middle of the interval. Interestingly, this effect of group size is already discernible, though not explained in the original study: in Figure 3 of Powers and Lehmann (2013), where group size is larger, *h* is also closer to 0.5 than in Figure 2 of Powers and Lehmann (2013) (middle panel).

##### A different social choice function leads to a more optimal institution

We then conduct the same analysis as before, this time varying the social choice function, replicating the two used in the original study (average and random dictator), and adding median and novelty. According to our predictions, this last social choice function should result in stronger selection, leading to the evolution of institutions that minimise waste of the public good. This is exactly what we observe: at equilibrium, the proportion of public good wasted is 10 times lower, around 2.5%. The mean value of the institution evolves to a much higher level, around 0.95 instead of 0.75, suggesting that the optimal value of the institution is higher than might have been assumed based on earlier results. This confirms that selection under novelty is stronger and thus more effective at improving institutions.

##### Limits to coevolution between institutions and demography

Examining different demographic scenarios also shows that increasing the carrying capacity has a diminishing effect on group size. For instance, while increasing the parameter *β* tenfold (from 3,000 to 30,000) roughly doubles average group size, further increasing it tenfold results in a barely perceivable increase in size. Our findings suggest a hypothesis to explain this pattern. As groups grow, selection weakens and waste of the public good increases, which counterbalances the effect of additional carrying capacity. In other words, the larger benefit produced by the public good leads to larger groups, but these larger groups experience weaker selection and waste a greater share of the public good. Ultimately, group size may not be limited by carrying capacity itself, but by the population’s ability to evolve optimal institutions. If the hypothesis is supported, this suggests that the coevolutionary process described in the original study between institutions and demography may operate only within a narrower range of conditions than initially thought—for example in medium-scale societies—or would require changes in how institutions are decided in order to persist.

#### Appendix E.2.4 Conclusion

In this section, we replicated and extended the model of Powers and Lehmann (2013) to test our framework in a richer, mechanistic setting. These results should not be read as direct predictions for real-world populations, nor as a critique of previous conclusions, which have been examined more fully in earlier work. Our implementation relies on a specific mutation process with strong drift, and exploring alternative assumptions would require a dedicated study beyond the scope of this paper. Our aim here is rather (i) to show how our findings clarify existing results and (ii) to test our predictions in more complex settings.

As such, we believe the results in this section serve both purposes. First, they show that the insights developed in this paper help explain the behaviour of this class of models—for example, why evolved institutions may display persistent inefficiencies, or why increasing carrying capacity has diminishing returns on group size. Second, they confirm that the key predictions of our analytical framework, concerning the effects of aggregation rules, group size, and selection strength, continue to hold when extended beyond abstract payoff functions.

